# AFA: Computationally efficient Ancestral Frequency estimation in Admixed populations: the Hispanic Community Health Study/Study of Latinos

**DOI:** 10.1101/2021.08.06.455462

**Authors:** Einat Granot-Hershkovitz, Quan Sun, Maria Argos, Hufeng Zhou, Xihong Lin, Sharon R. Browning, Tamar Sofer

## Abstract

We developed a computationally efficient method, Ancestral Frequency estimation in Admixed populations (AFA), to estimate the frequencies of bi-allelic variants in admixed populations with an unlimited number of ancestries. AFA uses maximum likelihood estimation by modeling the conditional probability of having an allele given proportions of genetic ancestries. It can be applied using either global or local proportions of genetic ancestries. Simulations mimicking admixture demonstrated the high accuracy of the method. We implemented the method on data from the Hispanic Community Health Study/Study of Latinos (HCHS/SOL), an admixed population with three predominant continental ancestries: Amerindian, European, and African. Comparison of the European and African estimated frequencies to the respective gnomAD frequencies demonstrated high correlations, with Pearson R^2^=0.97-0.99. We provide a genome-wide dataset of the estimated three ancestral allele frequencies in HCHS/SOL for all available variants with allele frequency between 5%-95% in at least one of the three ancestral populations.

## Introduction

Admixed populations have multiple ancestral origins, with different admixture patterns within and between populations, resulting from historical worldwide migration of populations^1^. Estimation of ancestry-specific allele frequencies in admixed populations can identify ancestry-specific enriched variants, with higher minor allele frequencies (MAFs) in one ancestry, compared to other ancestries. Fine mapping of association regions detected in admixture mapping, where one tests the association between local ancestry genomic interval (LAI) counts and a trait, can prioritize ancestry-specific enriched variants located in the identified regions for conditional association testing^2,3^. Similarly, genome-wide association studies (GWAS) of admixed populations can be followed by replication testing in homogeneous populations from a specific ancestry chosen based on the associated variant’s ancestry-specific frequencies. More generally, allele frequencies are important for interpreting sequence variants, distinguishing between pathogenic and benign variants^4^, inferring demographic histories of populations, and determining susceptibility to disease^5^. Thus, ancestry-specific allele frequencies can contribute to both research and personalized medicine of admixed populations. This is especially relevant for modern-day populations that are becoming increasingly genetically admixed^6^.

Several population genetic software packages were previously developed for admixture and population structure analyses, producing a by-product of ancestry-specific allele frequencies estimation in admixed populations^7,8^. Gravel et al. developed an algorithm based on the expectation-maximization (EM) framework relying on LAIs; but their method is not publicly available^9^. A similar publicly available algorithm, ASAFE, was developed. However, this method is available only for a three-way admixed diploid population, for genotyped markers located in LAIs, and it is time-consuming^10^. ASAFE was later extended to multi-way admixed populations in an algorithm that maximizes a multinomial likelihood^11^. Unfortunately, the software was not made public.

Here, we developed a computationally efficient method, Ancestral Frequency estimation in Admixed populations (AFA), for the estimation of ancestry-specific allele frequencies for bi-allelic variants, in a multi-way (unlimited) admixed population, with no need for phased data. Our model is similar to that proposed by Gravel et al., using maximum likelihood estimation by modeling the conditional probability of having a variant allele given local proportion ancestries (LAFA). We further extended the model by leveraging global ancestry proportions (GAFA), which are easier to compute and are more widely available, and we provide publicly available code. We examined the accuracy of our method by applying it to a simulated three-way admixed dataset. We then implemented the method on imputed genome-wide genetic data from the Hispanic Community Health Study/Study of Latinos (HCHS/SOL), an admixed population previously characterized with three predominant continental ancestries: Amerindian, European, and African, with varying proportions between individuals^12^. We computed ancestry-specific frequencies through AFA using the previous global proportion ancestries calculated by ADMIXTURE^12^ and LAIs calculated by RFMix^13,14^. We hypothesized that frequency estimates of variants using local ancestries (LAFA) would be more precise than estimates using global proportion ancestries (GAFA). We compared our estimated ancestral-specific variant frequencies for European and African ancestries to their respective frequencies published in gnomAD, expecting them to be similar, though not identical. Finally, we provide estimated Hispanic/Latino ancestry-specific allele frequencies estimated based on the HCHS/SOL for all variants with allele frequency between 5%-95% in at least one of the three ancestral populations.

## Methods

### Study population

The HCHS/SOL is a population-based longitudinal cohort study of US Hispanics/Latinos with participants recruited from four field centers (Bronx, NY, Chicago, IL, Miami, FL, and San Diego, CA) by a sampling design previously described ^15,16^. A total of 16,415 self-identified Hispanic/Latino adults, 18-to 74-year-old, were recruited during the first visit between 2008 and 2011, and various biospecimen and health information about risk/protective factors were collected.

### Genetic data

Genotyping and quality control were previously described ^12,17^. In brief, genotyping was performed using Illumina MEGA array, and a total of 11,928 samples and 985,405 genotyped variants passed quality control. Genome-wide imputation was conducted using the multi-ethnic NHLBI Trans-Omics for Precision Medicine (TOPMed) freeze 8 reference panel (GRCh38 assembly)^18^. Due to the overlap of samples in our target data and the TOPMed freeze 8 reference panel (n=6,201), we recalculated the estimated imputation quality (R2) using only non-overlapped samples to avoid over-estimates of the imputation quality. After filtering variants with R2<0.6 and minor allele count ≤5, a total of 42,038,818 imputed variants remained for analysis. Coordinates of genotyped and imputed variants were converted from GRCh38 to GRCh37 using the liftOver tool from UCSC^19^ for LAFA analysis since the LAIs were based on GRCh37 (as described below).

### Global proportion ancestries

Global continental ancestry proportions were previously estimated for 9,864 unrelated HCHS/SOL individuals using ADMIXTURE software under the assumption of three ancestral populations (Amerindian, African, and European), based on reference panels representing these ancestral populations^12^. After excluding individuals to generate a data set in which none of the individuals are third-degree relatives or closer, and individuals who withdrew consent for genetic studies, 8,933 individuals remained.

### Local ancestry intervals (LAIs)

Three-way LAI (Amerindian, African, and European) were previously inferred in 12,793 HCHS/SOL individuals using the RFMix software with a reference panel derived from the combination of the Human Genome Diversity Project (HGDP) and the 1000 Genome Project (using the GRCh37 assembly) representing the relevant ancestral populations^20^. Overall, 15,500 are LAIs dispersed throughout the genome (14,815 LAI in autosomal chromosomes), each spanning ten to hundreds of thousands of base pairs. After excluding individuals to generate a data set in which none of the individuals are third-degree relatives or closer, and individuals who withdrew consent for genetic studies, 9,512 individuals remained.

All participants in this analysis signed informed consent in their preferred language (Spanish/English) to use their genetic data. The study was reviewed and approved by the Institutional Review Boards at all collaborating institutions.

### Statistical analysis

#### The statistical model for estimation of ancestry-specific allele frequencies in admixed populations (AFA)

Suppose that we have a population of *n* individuals with *K* genetic ancestries. Consider a specific bi-allelic genetic variant in an autosomal chromosome. Each person has two copies of a variant potentially inherited from different ancestries. The genetic ancestry of each copy of the variant was inherited from the local ancestry encompassing the variant. For any given variant allele *g*, denote its ancestry-specific frequencies by *f*_1_, … , *f*_*K*_ in ancestries 1, … , *K*, respectively. Denote further the probability that person *i* has local ancestry *k* at the variant by *p*_*i,k*_, *k* = 1, … , *K*. We have that *p*_*i*,1_ … , *p*_*i,K*_ satisfy 0 ≤ *p*_*i,k*_ ≤ 1 and *p*_*i*,1_ + ⋯ + *p*_*i,K*_ = 1, for *i* = 1, … , *n, k* = 1, … , *K*. The allele count at the variant on a given chromosomal copy is sampled from a mixture of Bernoulli distributions, with

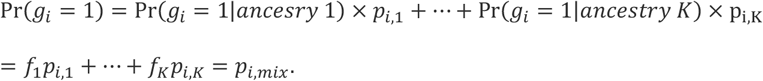

For unphased data, or when using genetic ancestry probabilities that are not specific to the variant (e.g., global ancestries), the probabilities *p*_*i*,1_, … , *p*_*i,K*_ are the same for the two copies of the allele. Under Hardy-Weinberg equilibrium at each ancestry, we can extend the model above to a Binomial distribution with two alleles. If *g*_*i*_ is now a bi-allelic variant, then:

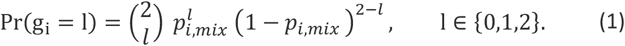

Assuming ancestral probabilities *p*_*i*,1_, … , *p*_*i,K*_ are known, the unknown frequencies *f*_1_, … , *f*_*K*_ can now be estimated by maximizing the log-likelihood across the sample of independent individuals. The standard errors of the estimated frequencies can be used to compute confidence intervals. We use the base R optim function with the “L-BFGS-B” optimization method for *K* > 1 ancestries and the “Brent” method when estimating allele frequency in one ancestry (for example, if *K* − 1 for *K* > 1 frequencies are known or assumed).

#### Choosing probabilities of genetic ancestry at the variant

To maximize the likelihood above, we assume that the ancestral probabilities *p*_*i*,1_, … , *p*_*i,K*_ of the study individuals are known. In practice, they are estimated. We consider two estimators. First is the global proportion of ancestry (GAFA). These could be computed using software packages such as ADMIXTURE or RFMix, with a subset of independent, genotyped genetic variants, with or without a reference panel^6–8,14^. The second estimator is based on LAIs (LAFA). Local ancestry analysis results in a segmentation of the genome in which each segment, LAI, is assigned a genetic ancestry. Thus, a given variant *g* is overlapping with a certain LAI, say *LAI*_*g*_, which is annotated with two genetic ancestries. With some local ancestry inference methods, such as RFMix, these LAIs are unphased with respect to the allele counts. To generate a vector of genetic ancestry probabilities for the variant, we first generate a vector of counts of local ancestries (*c*_*i*,1_, … , *c*_*i,K*_) , and divide all entries by two, the highest attained count. In mathematical notation:

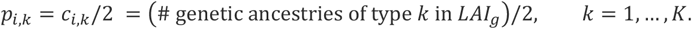

The probabilities here take values 0, 0.5, 1.

#### Computing ancestry-specific allele frequencies on the X chromosome

The methodology for the X-chromosome is similar, with a slight difference for males, where we use a Bernoulli distribution (or a Binomial distribution with parameters (*p*_*mix*_, 1)) to account for the fact that there is a single observed allele.

#### Handling of boundary conditions

The log-likelihood of the Binomial distribution cannot be maximized at the boundaries, i.e. when the data is consistent with an ancestry-specific frequency at the boundary of the parameter space, e.g. *f*_*k*_ ∈ {0,1} for some *k* = 1, … , *K* . To prevent non-convergence of the estimation algorithm, we implemented a procedure that generates synthetic observations and adds them to the data. These are 2*K* synthetic observations, two for each ancestry, mimicking a reference and alternate allele from each of the genetic ancestries. For example, one synthetic observation will have a single reference allele for a (simulated) person, and the ancestral probabilities for this person are *p*_*i,k*_ = 1 for ancestry *k*, and *p*_*i,l*_ = 0 for all other ancestries *l* ≠ *k, l* ∈ {1, … , *K*}. Another synthetic observation will have a single alternate allele for this variant, and the same values of ancestral probabilities. In addition, the algorithm allows for settings box constraints on the boundaries^21^.

#### An approximation for computing ancestry-specific allele frequency using imputed data

When imputed data are confidently estimated, the extension of the algorithm to imputed genotypes is straightforward. For imputed genotypes with fractions, we cannot compute the log-likelihood based on the probability in (1). Instead, we notice that we can decompose the function into two parts: “2 choose *l*” and 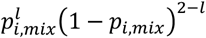. The second part can be computed for any *l*, while the first part cannot. Instead, we apply linear interpolation to compute a value approximating “2 choose *l*” based on the values of this function evaluated at the nearest integers higher and lower than *l*.

#### Simulation studies

We studied our method, AFA, in simulations to determine how the ancestral frequency estimation accuracy is influenced by the effective sample size, effn, defined as 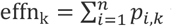 for ancestry *k* = 1, … , *K*, by the expected allele frequencies (rare vs. common variants) and by using the local vs. global proportion ancestries (LAFA vs. GAFA).

We simulated a three-way admixed population, using fixed effn_1_=effn_2_=1,000, varied effn_3_ in the range 100-4,000, and focused on the estimation of *f*_3_. We fixed *f*_1_ = 0.5, *f*_2_ = 0.3 throughout, and varied the allele frequency *f*_3_ ∈ {0.01,0.05,0.1,0.2}. First, we simulated local ancestries based on global effn (where n=effn_1_ + effn_2_ + effn_3_). We assumed that each person has two copies of 10 LAIs of equal lengths. Thus, the overall number of LAIs of ancestry *k* ∈ {1, 2,3} was *n* × effn_k_ ∗ 20. Then, we randomly assigned 20 LAIs (2 copies of 10 LAIs) to individuals and computed the global proportion of ancestries for each individual as the proportion of LAIs of each ancestry. The genetic variant was assumed to be in the first LAI. Next, we simulated the allele counts based on the allele frequencies *f*_1_, *f*_2_, and *f*_3_. For each person and each copy of the first LAI, we sampled the allele from the Bernoulli distribution with a probability according to the ancestry at the interval copy. To mimic the real data, which is unphased, we then summed the allele count across the two copies for each person. Finally, we estimated ancestry-specific allele frequencies using the computed global proportion ancestries and using the ancestries of the first LAI. We performed *n*_*sim*_ =1,000 simulation replicates for each setting. We also performed a similar simulation based on a homogenous population derived from a single ancestry to compare the expected bias in frequency estimation in admixed populations to that in a non-Admixed population when using the same algorithm.

Let 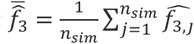 denote the mean estimated *f*_3_ across simulations. We assessed the frequency estimation accuracy of *f*_3_ using the following measures:

1. Bias: 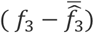.
2. Inflation: 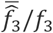.
3. RMSE (root mean squared error): 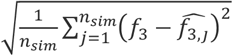

#### Comparing ancestry-specific allele frequency estimates to previously published estimates

We compared the estimated ancestry-specific frequencies of 9 variants using GAFA and LAFA, with previously published estimated ancestral frequencies based on the ASAFE method in the HCHS/SOL dataset^3,22,23^. We also compared the estimated Amerindian frequency of 4 variants with the previously published frequencies in Pima-Indians^3^.

#### Comparing estimated ancestry-specific allele frequencies to gnomAD allele frequencies

We compared the estimated European and African frequencies in the admixed HCHS/SOL population using GAFA and LAFA to the gnomAD v2 liftover (GRCh38) non-Finnish European and African frequencies, respectively, by plotting and calculating the Pearson squared correlation coefficient. We assessed only gnomAD variants passing quality control filters (FILTER==“PASS”), with an ancestral minor allele count of ≥100 respective to the assessed ancestry. We also calculated the percentage of estimated confidence intervals (CIs) for ancestral MAFs using GAFA and LAFA, which include the concordant reported gnomAD MAFs, binned by gnomAD MAF categories.

#### Availability and implementation

We provide a publicly available GitHub repository, https://github.com/tamartsi/Ancestry_specific_freqs, which includes: (1) code for GAFA and LAFA for computing ancestry-specific allele frequencies (2) simulation code (3) a dataset of Hispanic/Latino ancestry-specific allele frequencies and their Cls estimated based on the HCHS/SOL using GAFA and LAFA for all variants (genotyped or imputed) with an estimated frequency between 5%-95% in at least one of the three ancestral populations. This dataset will also be available through FAVOR (Functional Annotation of Variants – Online Resource) v2 data release in both the single variant query (Allele Frequency Block) and batch query, http://favor.genohub.org. CWL workflows for GAFA and LAFA are also available via dockstore and https://github.com/cwl-apps/ancestral-maf-admixed-population.

## Results

### Simulation studies

Table 1 and Figure 1 summarize the results from simulation studies of frequency estimation in a three-way admixed population, based on GAFA or LAFA. For comparison, simulation results based on non-admixed populations under the same framework, essentially reducing to standard maximum likelihood estimation, are presented in Supplementary Table 1 and Supplementary Figure 1. As expected, estimated frequencies become more accurate with increasing effective sample size and increasing MAF. Likely due to the boundaries of the parameter space, the estimated frequencies tend to be biased towards more common MAFs until large enough effective sample sizes or allele frequencies (or in other words, enough counts of the minor allele) are available. In addition, accuracy increased when using LAFA compared to GAFA. For example, for MAF =0.01 and effn=4,000 we had bias =0.00574 for GAFA and bias= 0.00022 for LAFA; for MAF=0.2 and effn=1000 we had bias = 0.00453 for GAFA and bias = 0.00028 for LAFA (Table1). Similar trends of improved accuracy of frequency estimation with larger effective sample sizes and higher MAFs are observed in the non-admixed population analysis (Supplementary Table 1 and Supplementary Figure 1), with, unsurprisingly, higher accuracy compared to the admixed population.

**Table 1:** Results from simulation studies of frequency estimation of a bi-allelic variant in a three-way admixed population, based on AFA (Ancestral Frequency estimation in Admixed populations), by different effective sample sizes and different expected minor allele frequencies. For each of the settings, we tested 1,000 simulation replicates and calculated the mean frequency estimate, the difference, and ratio of the mean observed frequency and the expected frequency, the RMSE of the estimate frequncies, the percentage of CI including the expected frequency (coverage), and the 95% interval of the estimated frequencies. The results refer to one of the ancestries. The characteristics of the other two ancestries were the same in all simulations, with effective sample size of effn=1,000, one ancestry with MAF=0.5 and the other with MAF=0.3.

| Expect<br>ed | Effective<br>N | Three-way admixed population using GAFA |  |  |  |  |  |  | Three-way admixed population using LAFA |  |  |  |  |  |  |
| --- | --- | --- | --- | --- | --- | --- | --- | --- | --- | --- | --- | --- | --- | --- | --- |
|  |  | MAF<br>Mean | MAF<br>difference | MAF<br>ratio | RMSE | CI %<br>coverage | interval<br>2.5% | interval<br>97.5% | MAF<br>Mean | MAF<br>difference | MAF<br>ratio | RMSE | CI %<br>coverage | interval<br>2.5% | interval<br>97.5% |
| 0.005 | 100 | 0.153 | 0.148 | 30.556 | 0.164 | 0.90 | 0.0569 | 0.3284 | 0.040 | 0.035 | 8.001 | 0.039 | 0.97 | 0.0172 | 0.0797 |
|  | 200 | 0.109 | 0.104 | 21.736 | 0.115 | 0.90 | 0.0407 | 0.2264 | 0.023 | 0.018 | 4.643 | 0.021 | 0.98 | 0.0102 | 0.0499 |
|  | 500 | 0.063 | 0.058 | 12.683 | 0.065 | 0.90 | 0.0251 | 0.1327 | 0.009 | 0.004 | 1.856 | 0.006 | 0.99 | 0.0038 | 0.0194 |
|  | 1000 | 0.040 | 0.035 | 7.981 | 0.040 | 0.91 | 0.0145 | 0.0866 | 0.007 | 0.002 | 1.380 | 0.003 | 0.99 | 0.0029 | 0.0123 |
|  | 2000 | 0.023 | 0.018 | 4.542 | 0.020 | 0.93 | 0.0088 | 0.0471 | 0.006 | 0.001 | 1.105 | 0.002 | 0.94 | 0.0027 | 0.0091 |
|  | 4000 | 0.012 | 0.007 | 2.468 | 0.009 | 0.93 | 0.0047 | 0.0253 | 0.005 | 0.000 | 1.060 | 0.001 | 0.94 | 0.0035 | 0.0072 |
| 0.01 | 100 | 0.156 | 0.146 | 15.631 | 0.164 | 0.90 | 0.0609 | 0.3369 | 0.043 | 0.033 | 4.306 | 0.038 | 0.97 | 0.0174 | 0.0865 |
|  | 200 | 0.110 | 0.100 | 11.013 | 0.112 | 0.91 | 0.0424 | 0.2340 | 0.027 | 0.017 | 2.651 | 0.020 | 0.98 | 0.0103 | 0.0563 |
|  | 500 | 0.066 | 0.056 | 6.645 | 0.064 | 0.92 | 0.0247 | 0.1395 | 0.014 | 0.004 | 1.401 | 0.007 | 0.99 | 0.0043 | 0.0272 |
|  | 1000 | 0.044 | 0.034 | 4.383 | 0.039 | 0.91 | 0.0171 | 0.0928 | 0.012 | 0.002 | 1.156 | 0.004 | 0.96 | 0.0050 | 0.0195 |
|  | 2000 | 0.026 | 0.016 | 2.622 | 0.020 | 0.93 | 0.0093 | 0.0562 | 0.011 | 0.001 | 1.053 | 0.002 | 0.95 | 0.0067 | 0.0151 |
|  | 4000 | 0.016 | 0.006 | 1.574 | 0.009 | 0.95 | 0.0061 | 0.0303 | 0.010 | 0.000 | 1.022 | 0.001 | 0.94 | 0.0074 | 0.0129 |
| 0.05 | 100 | 0.176 | 0.126 | 3.524 | 0.149 | 0.94 | 0.0685 | 0.3589 | 0.071 | 0.021 | 1.410 | 0.034 | 0.97 | 0.0271 | 0.1327 |
|  | 200 | 0.134 | 0.084 | 2.677 | 0.102 | 0.94 | 0.0500 | 0.2681 | 0.060 | 0.010 | 1.208 | 0.024 | 0.95 | 0.0235 | 0.1071 |
|  | 500 | 0.091 | 0.041 | 1.810 | 0.056 | 0.94 | 0.0344 | 0.1757 | 0.053 | 0.003 | 1.063 | 0.013 | 0.93 | 0.0290 | 0.0797 |
|  | 1000 | 0.069 | 0.019 | 1.388 | 0.033 | 0.95 | 0.0259 | 0.1258 | 0.051 | 0.001 | 1.020 | 0.008 | 0.96 | 0.0373 | 0.0660 |
|  | 2000 | 0.056 | 0.006 | 1.117 | 0.018 | 0.97 | 0.0265 | 0.0887 | 0.050 | 0.000 | 1.002 | 0.004 | 0.96 | 0.0413 | 0.0588 |
|  | 4000 | 0.052 | 0.002 | 1.035 | 0.010 | 0.95 | 0.0336 | 0.0716 | 0.050 | 0.000 | 1.004 | 0.003 | 0.95 | 0.0446 | 0.0561 |
| 0.1 | 100 | 0.200 | 0.100 | 2.003 | 0.134 | 0.95 | 0.0744 | 0.4059 | 0.115 | 0.015 | 1.152 | 0.038 | 0.96 | 0.0512 | 0.1863 |
|  | 200 | 0.164 | 0.064 | 1.641 | 0.093 | 0.94 | 0.0611 | 0.3204 | 0.107 | 0.007 | 1.068 | 0.026 | 0.96 | 0.0627 | 0.1587 |
|  | 500 | 0.126 | 0.026 | 1.259 | 0.052 | 0.95 | 0.0504 | 0.2293 | 0.102 | 0.002 | 1.023 | 0.015 | 0.96 | 0.0742 | 0.1320 |
|  | 1000 | 0.110 | 0.010 | 1.104 | 0.033 | 0.95 | 0.0512 | 0.1744 | 0.101 | 0.001 | 1.009 | 0.010 | 0.95 | 0.0823 | 0.1214 |
|  | 2000 | 0.103 | 0.003 | 1.031 | 0.021 | 0.95 | 0.0640 | 0.1433 | 0.100 | 0.000 | 1.001 | 0.006 | 0.95 | 0.0882 | 0.1131 |
|  | 4000 | 0.101 | 0.001 | 1.009 | 0.011 | 0.97 | 0.0796 | 0.1217 | 0.100 | 0.000 | 1.003 | 0.004 | 0.95 | 0.0929 | 0.1084 |
| 0.2 | 100 | 0.264 | 0.064 | 1.322 | 0.123 | 0.96 | 0.0984 | 0.4848 | 0.209 | 0.009 | 1.044 | 0.043 | 0.96 | 0.1275 | 0.2906 |
|  | 200 | 0.234 | 0.034 | 1.171 | 0.086 | 0.95 | 0.0955 | 0.4126 | 0.203 | 0.003 | 1.015 | 0.029 | 0.96 | 0.1466 | 0.2588 |
|  | 500 | 0.214 | 0.014 | 1.069 | 0.059 | 0.95 | 0.1062 | 0.3281 | 0.201 | 0.001 | 1.006 | 0.018 | 0.94 | 0.1655 | 0.2379 |
|  | 1000 | 0.205 | 0.005 | 1.023 | 0.036 | 0.96 | 0.1351 | 0.2749 | 0.200 | 0.000 | 1.001 | 0.012 | 0.96 | 0.1764 | 0.2217 |
|  | 2000 | 0.201 | 0.001 | 1.005 | 0.023 | 0.95 | 0.1571 | 0.2468 | 0.200 | 0.000 | 1.002 | 0.008 | 0.95 | 0.1844 | 0.2152 |
|  | 4000 | 0.200 | 0.000 | 1.000 | 0.013 | 0.96 | 0.1755 | 0.2240 | 0.200 | 0.000 | 1.000 | 0.005 | 0.95 | 0.1895 | 0.2103 |

Abbreviations: *GAFA* Global -Ancestral Frequency estimation in Admixed populations; *LAFA* Local -Ancestral Frequency estimation in Admixed populations; *MAF* minor allele frequency; *CI* confidence interval; *RMSE* root mean squared error.

**Figure 1:**
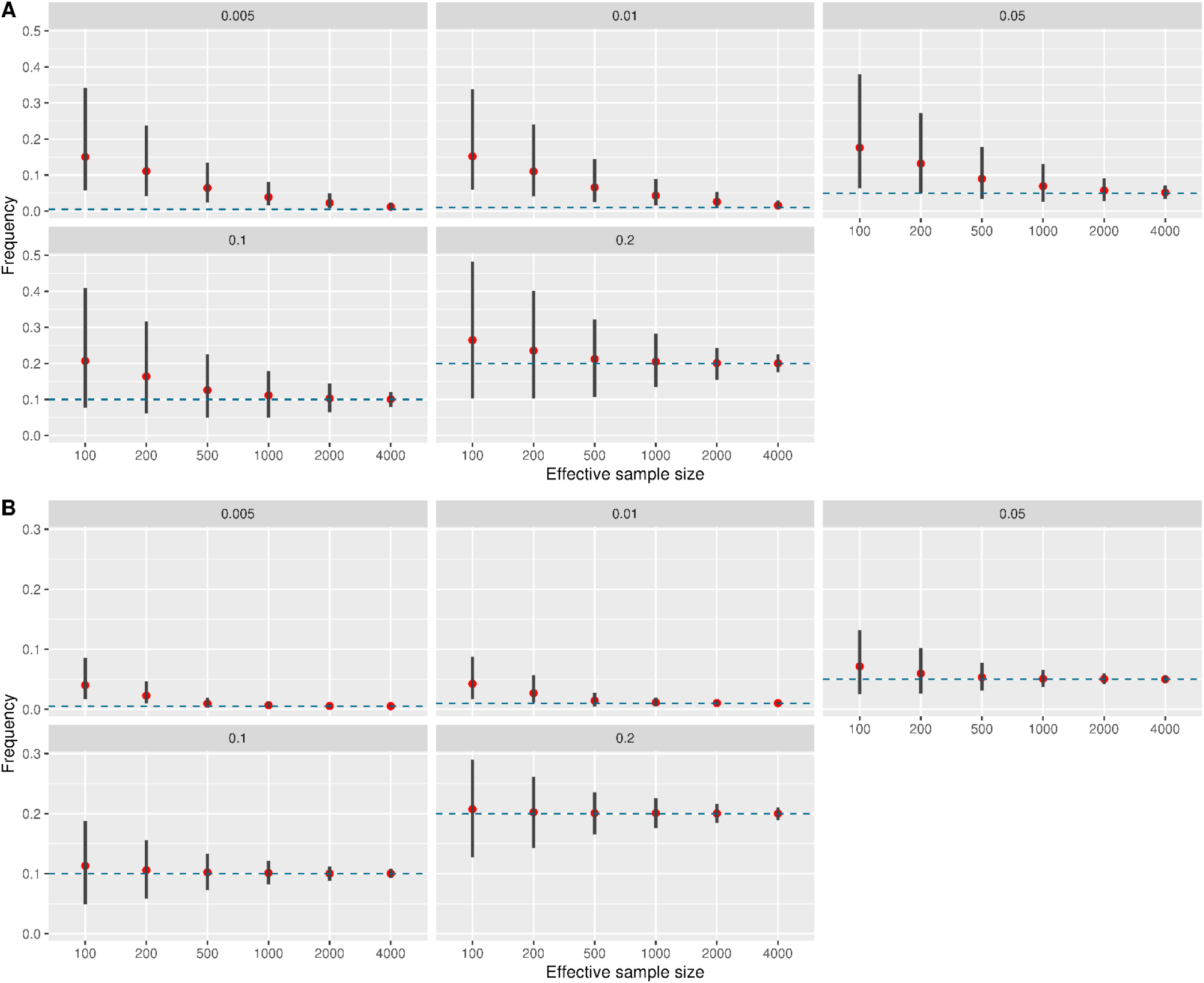
Results from simulation studies of frequency estimation of a bi-allelic variant in a three-way admixed population, based on A. GAFA (Global -Ancestral Frequency estimation in Admixed populations) B. LAFA (Local -Ancestral Frequency estimation in Admixed populations). Various settings include a different effective sample size of effn (x-axis) and different expected minor allele frequencies (indicated in the upper title of each graph). We performed 1,000 simulation replicates of each scenario. Each dot represents the mean frequency of 1,000 simulation replicates each line represents the 95% interval estimated frequencies across the simulation replicates.

### Hispanic Community Health Study/Study of Latinos

We applied AFA to the HCHS/SOL imputed dataset, excluding variants with minor allele count ≤5, setting frequency boundary conditions (low= 0.00001, high = 0.99999) as arguments to the optimization function. If AFA did not converge for a given variant, we applied it again with a stricter boundary condition (low=0.01, high =0.99). We developed workflows for GAFA and LAFA on BioData Catalyst Powered by Seven Bridges (https://biodatacatalyst.nhlbi.nih.gov/). We processed data in a parallel manner by batching the workflows by chromosomes and scattering jobs by blocks of 3,000 variants, using the c5.18xlarge spot instance provisioned on Amazon Web Services. The workflows are described (represented) in the Common Workflow Language open standard^24^ and are therefore portable to multiple computational environments. The computation time for the shortest chromosome (chr21, n=552,556 variants) was 57 minutes using GAFA and 110 minutes using LAFA, with ∼50 jobs running in parallel. The number of estimated variant frequencies per chromosome is summarized in Supplementary Table 2 stratified by boundary condition, for both GAFA and LAFA. The number of variants for which we provide estimated variant frequencies, under the condition that they have a frequency between 5%-95% in at least one of the three ancestral populations, is summarized in Table 2 stratified by boundary condition, for both GAFA and LAFA. In general, rare variants required strict boundary conditions (0.01 rather than 0.00001) on the estimated frequencies for algorithm convergence.

**Table 2:** Number of estimated variant frequencies per chromosome in HCHS/SOL that are common (frequency between 5%-95%) in at least one of the three ancestral populations, stratified by boundary condition, calculated via GAFA or LAFA.

| Chromosome | GAFA |  |  | LAFA |  |  |
| --- | --- | --- | --- | --- | --- | --- |
|  | Total | Boundary 1E-05 (%) | Boundary 1E-02 | Total | Boundary 1E-05 (%) | Boundary 1E-02 |
| 1 | 731372 | 583,809 (79.82) | 147563 | 733709 | 413,042 (56.3) | 320667 |
| 2 | 789792 | 629,106 (79.65) | 160686 | 812360 | 448,927 (55.26) | 363433 |
| 3 | 683573 | 547,793 (80.14) | 135780 | 692240 | 389,055 (56.2) | 303185 |
| 4 | 699617 | 560,836 (80.16) | 138781 | 703178 | 407,366 (57.93) | 295812 |
| 5 | 608090 | 488,543 (80.34) | 119547 | 622061 | 343,828 (55.27) | 278233 |
| 6 | 623354 | 516,241 (82.82) | 107113 | 625767 | 355,100 (56.75) | 270667 |
| 7 | 557900 | 455,514 (81.65) | 102386 | 557945 | 321,508 (57.62) | 236437 |
| 8 | 530427 | 419,391 (79.07) | 111036 | 545404 | 294,384 (53.98) | 251020 |
| 9 | 411577 | 332,874 (80.88) | 78703 | 417816 | 232,124 (55.56) | 185692 |
| 10 | 483953 | 394,154 (81.44) | 89799 | 481662 | 276,438 (57.39) | 205224 |
| 11 | 470444 | 378,998 (80.56) | 91446 | 479053 | 269,368 (56.23) | 209685 |
| 12 | 458340 | 367,421 (80.16) | 90919 | 459471 | 257,015 (55.94) | 202456 |
| 13 | 348128 | 288,100 (82.76) | 60028 | 353459 | 208,717 (59.05) | 144742 |
| 14 | 307247 | 250,059 (81.39) | 57188 | 307967 | 171,930 (55.83) | 136037 |
| 15 | 271244 | 219,250 (80.83) | 51994 | 275865 | 155,716 (56.45) | 120149 |
| 16 | 285290 | 226,155 (79.27) | 59135 | 279954 | 158,444 (56.6) | 121510 |
| 17 | 256758 | 207,176 (80.69) | 49582 | 250340 | 140,177 (55.99) | 110163 |
| 18 | 270605 | 221,435 (81.83) | 49170 | 273281 | 158,435 (57.98) | 114846 |
| 19 | 215685 | 176,195 (81.69) | 39490 | 202739 | 115,992 (57.21) | 86747 |
| 20 | 212499 | 169,496 (79.76) | 43003 | 212818 | 118,307 (55.59) | 94511 |
| 21 | 129185 | 105,026 (81.3) | 24159 | 131628 | 75,669 (57.49) | 55959 |
| 22 | 127717 | 104,683 (81.96) | 23034 | 123688 | 69,062 (55.84) | 54626 |
| X | 335292 | 235,553 (70.25) | 99739 | 301688 | 139,741 (46.32) | 161947 |
| Total | 9,808,089 | 7,877,808 (80.32) | 1,930,281 | 9,844,093 | 5,520,345 (56.08) | 4,323,748 |

Abbreviations: GAFA Global -Ancestral Frequency estimation in Admixed populations; LAFA Local -Ancestral Frequency estimation in Admixed populations.

### Comparing ancestry-specific allele frequency estimates to previously published estimates

Table 3 summarizes 9 previously published HCHS/SOL ancestry-specific allele frequencies estimated by ASAFE, for comparison with our GAFA and LAFA frequency estimations. Frequency estimations for all 9 variants are highly comparable, with absolute mean frequency differences for African=0.0008 European=0.0153 and Amerindian=0.0101 for GAFA and African=0.0023 European=0.019 and Amerindian=0.0094 for LAFA. Table 4 summarizes 4 previously published allele frequencies of Pima-Indians to the Amerindian ancestral frequency estimated in HCHS/SOL based on GAFA and LAFA. Here too, the absolute mean frequency differences are low with GAFA=0.03 and LAFA=0.01.

**Table 3:**
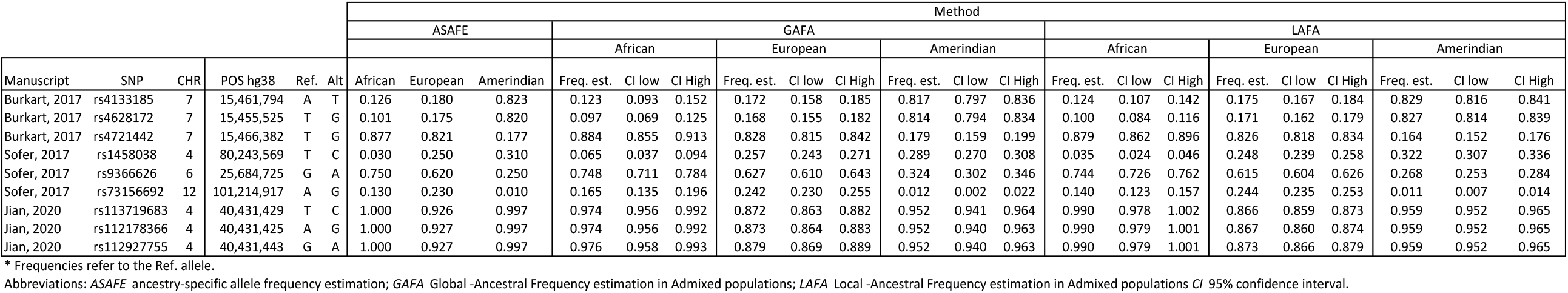
HCHS/SOL ancestry-specific allele frequencies previously published (estimated by ASAFE) compared to GAFA and LAFA frequency estimations.

|  |  |  |  |  |  | Method |  |  |  |  |  |  |  |  |  |  |  |  |  |  |  |  |  |  |  |  |
| --- | --- | --- | --- | --- | --- | --- | --- | --- | --- | --- | --- | --- | --- | --- | --- | --- | --- | --- | --- | --- | --- | --- | --- | --- | --- | --- |
|  |  |  |  |  |  | ASAFE |  |  | GAFA |  |  |  |  |  |  |  |  | LAFA |  |  |  |  |  |  |  |  |
|  |  |  |  |  |  |  |  |  | African |  |  | European |  |  | Amerindian |  |  | African |  |  | European |  |  | Amerindian |  |  |
| Manuscript | SNP | CHR | POS hg38 | Ref. | Alt | African | European | Amerindian | Freq. est. | CI low | CI High | Freq. est. | CI low | CI High | Freq. est. | CI low | CI High | Freq. est. | CI low | CI High | Freq. est. | CI low | CI High | Freq. est. | CI low | CI High |
| Burkart, 2017 | rs4133185 | 7 | 15,461,794 | A | T | 0.126 | 0.180 | 0.823 | 0.123 | 0.093 | 0.152 | 0.172 | 0.158 | 0.185 | 0.817 | 0.797 | 0.836 | 0.124 | 0.107 | 0.142 | 0.175 | 0.167 | 0.184 | 0.829 | 0.816 | 0.841 |
| Burkart, 2017 | rs4628172 | 7 | 15,455,525 | T | G | 0.101 | 0.175 | 0.820 | 0.097 | 0.069 | 0.125 | 0.168 | 0.155 | 0.182 | 0.814 | 0.794 | 0.834 | 0.100 | 0.084 | 0.116 | 0.171 | 0.162 | 0.179 | 0.827 | 0.814 | 0.839 |
| Burkart, 2017 | rs4721442 | 7 | 15,466,382 | T | G | 0.877 | 0.821 | 0.177 | 0.884 | 0.855 | 0.913 | 0.828 | 0.815 | 0.842 | 0.179 | 0.159 | 0.199 | 0.879 | 0.862 | 0.896 | 0.826 | 0.818 | 0.834 | 0.164 | 0.152 | 0.176 |
| Sofer, 2017 | rs1458038 | 4 | 80,243,569 | T | C | 0.030 | 0.250 | 0.310 | 0.065 | 0.037 | 0.094 | 0.257 | 0.243 | 0.271 | 0.289 | 0.270 | 0.308 | 0.035 | 0.024 | 0.046 | 0.248 | 0.239 | 0.258 | 0.322 | 0.307 | 0.336 |
| Sofer, 2017 | rs9366626 | 6 | 25,684,725 | G | A | 0.750 | 0.620 | 0.250 | 0.748 | 0.711 | 0.784 | 0.627 | 0.610 | 0.643 | 0.324 | 0.302 | 0.346 | 0.744 | 0.726 | 0.762 | 0.615 | 0.604 | 0.626 | 0.268 | 0.253 | 0.284 |
| Sofer, 2017 | rs73156692 | 12 | 101,214,917 | A | G | 0.130 | 0.230 | 0.010 | 0.165 | 0.135 | 0.196 | 0.242 | 0.230 | 0.255 | 0.012 | 0.002 | 0.022 | 0.140 | 0.123 | 0.157 | 0.244 | 0.235 | 0.253 | 0.011 | 0.007 | 0.014 |
| Jian, 2020 | rs113719683 | 4 | 40,431,429 | T | C | 1.000 | 0.926 | 0.997 | 0.974 | 0.956 | 0.992 | 0.872 | 0.863 | 0.882 | 0.952 | 0.941 | 0.964 | 0.990 | 0.978 | 1.002 | 0.866 | 0.859 | 0.873 | 0.959 | 0.952 | 0.965 |
| Jian, 2020 | rs112178366 | 4 | 40,431,425 | A | G | 1.000 | 0.927 | 0.997 | 0.974 | 0.956 | 0.992 | 0.873 | 0.864 | 0.883 | 0.952 | 0.940 | 0.963 | 0.990 | 0.979 | 1.001 | 0.867 | 0.860 | 0.874 | 0.959 | 0.952 | 0.965 |
| Jian, 2020 | rs112927755 | 4 | 40,431,443 | G | A | 1.000 | 0.927 | 0.997 | 0.976 | 0.958 | 0.993 | 0.879 | 0.869 | 0.889 | 0.952 | 0.940 | 0.963 | 0.990 | 0.979 | 1.001 | 0.873 | 0.866 | 0.879 | 0.959 | 0.952 | 0.965 |

\* Frequencies refer to the Ref. allele. Abbreviations: *ASAFE* ancestry-specific allele frequency estimation; *GAFA* Global -Ancestral Frequency estimation in Admixed populations; *LAFA* Local -Ancestral Frequency estimation in Admixed populations *CI* 95% confidence interval.

**Table 4:** Previously published Pima Indians allele frequencies, compared to our GAFA and LAFA Amerindian frequency estimations in the HCHS/SOL.

|  |  |  |  |  |  | Pima Indians | GAFA |  |  | LAFA |  |  |
| --- | --- | --- | --- | --- | --- | --- | --- | --- | --- | --- | --- | --- |
| Paper | SNP | CHR | POS hg38 | Ref. | Alt |  | Freq. est. | CI low | CI High | Freq. est. | CI low | CI High |
| Sofer, 2017 | rs75432840 | 6 | 34,143,031 | C | G | 0.28 | 0.398 | 0.379 | 0.417 | 0.287 | 0.272 | 0.303 |
| Sofer, 2017 | rs138977532 | 6 | 36,382,025 | C | T | 0.59 | 0.635 | 0.620 | 0.650 | 0.587 | 0.572 | 0.602 |
| Sofer, 2017 | rs139139046 | 11 | 71,452,308 | G | C | 0.87 | 0.831 | 0.819 | 0.844 | 0.823 | 0.813 | 0.833 |
| Sofer, 2017 | rs72849841 | 17 | 80,298,494 | C | T | 1.00 | 0.987 | 0.978 | 0.996 | 0.996 | 0.994 | 0.998 |

\* Frequencies refer to the Ref. allele. Abbreviations: *GAFA* Global -Ancestral Frequency estimation in Admixed populations; *LAFA* Local -Ancestral Frequency estimation in Admixed populations.

### Comparing estimated ancestry-specific allele frequencies to gnomAD allele frequencies

Figure 2 compares the estimated European- and African-specific allele frequencies in HCHS/SOL for variants on chromosome 2 using GAFA and LAFA to the gnomAD non-Finnish European and African frequencies, respectively. All other chromosomes’ comparisons are presented in Supplementary Figures 2 (GAFA) and 3 (LAFA). All estimated frequencies were highly correlated, with Pearson R^2^=0.97-0.99. We further calculated the percentage of ancestral gnomAD frequencies which are included in the corresponding CI estimated in HCHS/SOL by GAFA or LAFA, binned by gnomAD frequency categories (Table 5). The mean range of CIs was also calculated for each category and was consistently smaller for LAFA compared to GAFA since the ancestral determination for each variant is more accurate when using LAIs. Thus, LAFA resulted in a lower percentage of included gnomAD allele frequencies relative to GAFA; however, this does not indicate a superiority of GAFA over LAFA because of potentially true differences in ancestral frequencies in HCHS/SOL compared to gnomAD. The mean ranges of CIs are lower in low-frequency variant bins compared to the common frequency bins, both for GAFA and LAFA.

**Table 5:**
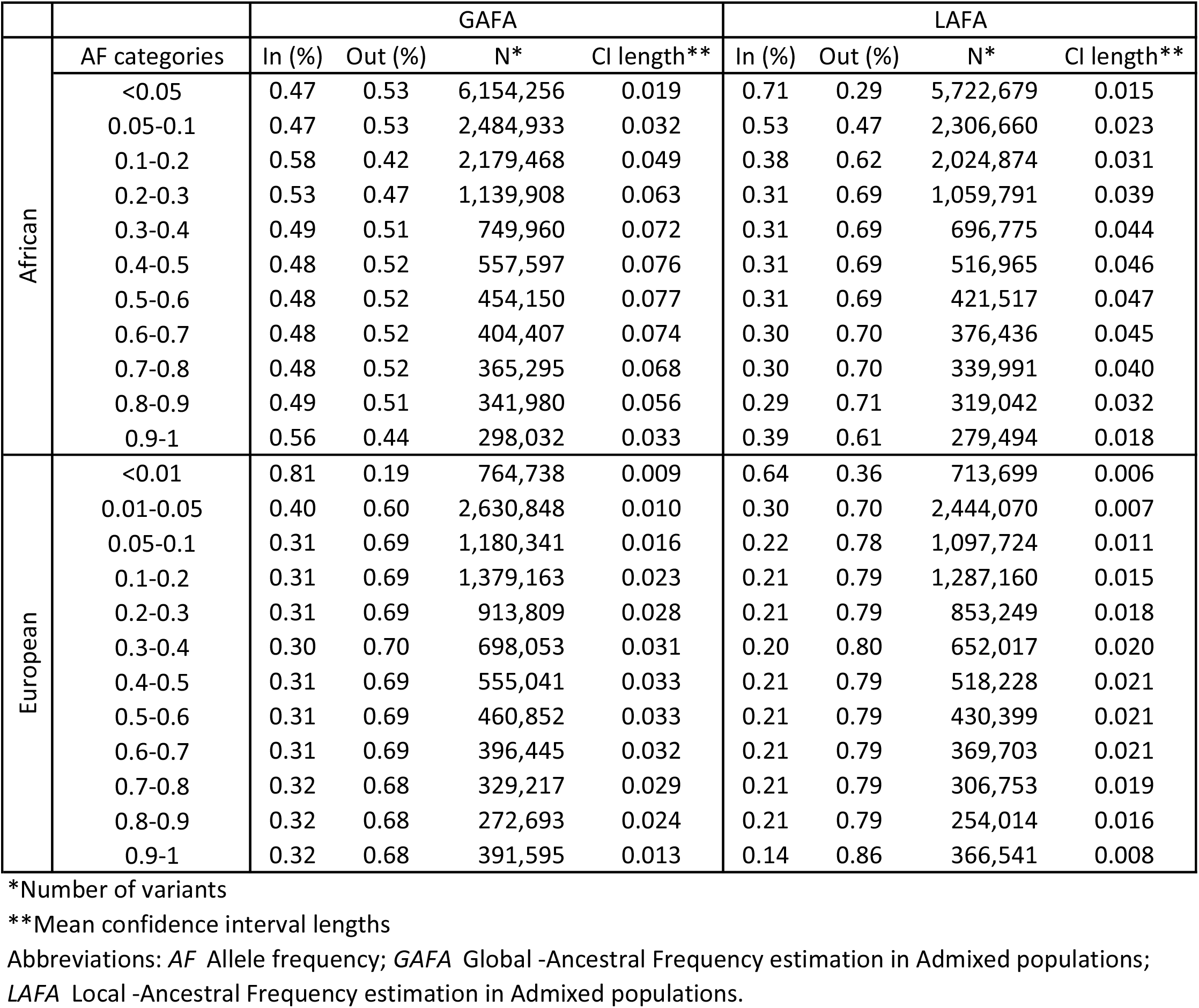
Percentage of non-Finnish European and African gnomAD frequencies included in the corresponding confidence interval (CI) estimated in HCHS/SOL by GAFA and LAFA, binned by gnomAD frequency categories. We assessed only gnomAD variants passing quality control filters (FILTER==“PASS”), with an ancestral minor allele count of ≥100 respective to the assessed ancestry.The Europeans have an extra category for rare variants (<0.01), since their calculation is based on a larger datset compared to Africans.

|  |  | GAFA |  |  |  | LAFA |  |  |  |
| --- | --- | --- | --- | --- | --- | --- | --- | --- | --- |
|  | AF categories | In (%) | Out (%) | N* | CI length** | In (%) | Out (%) | N* | CI length** |
| African | <0.05 | 0.47 | 0.53 | 6,154,256 | 0.019 | 0.71 | 0.29 | 5,722,679 | 0.015 |
|  | 0.05-0.1 | 0.47 | 0.53 | 2,484,933 | 0.032 | 0.53 | 0.47 | 2,306,660 | 0.023 |
|  | 0.1-0.2 | 0.58 | 0.42 | 2,179,468 | 0.049 | 0.38 | 0.62 | 2,024,874 | 0.031 |
|  | 0.2-0.3 | 0.53 | 0.47 | 1,139,908 | 0.063 | 0.31 | 0.69 | 1,059,791 | 0.039 |
|  | 0.3-0.4 | 0.49 | 0.51 | 749,960 | 0.072 | 0.31 | 0.69 | 696,775 | 0.044 |
|  | 0.4-0.5 | 0.48 | 0.52 | 557,597 | 0.076 | 0.31 | 0.69 | 516,965 | 0.046 |
|  | 0.5-0.6 | 0.48 | 0.52 | 454,150 | 0.077 | 0.31 | 0.69 | 421,517 | 0.047 |
|  | 0.6-0.7 | 0.48 | 0.52 | 404,407 | 0.074 | 0.30 | 0.70 | 376,436 | 0.045 |
|  | 0.7-0.8 | 0.48 | 0.52 | 365,295 | 0.068 | 0.30 | 0.70 | 339,991 | 0.040 |
|  | 0.8-0.9 | 0.49 | 0.51 | 341,980 | 0.056 | 0.29 | 0.71 | 319,042 | 0.032 |
|  | 0.9-1 | 0.56 | 0.44 | 298,032 | 0.033 | 0.39 | 0.61 | 279,494 | 0.018 |
| European | <0.01 | 0.81 | 0.19 | 764,738 | 0.009 | 0.64 | 0.36 | 713,699 | 0.006 |
|  | 0.01-0.05 | 0.40 | 0.60 | 2,630,848 | 0.010 | 0.30 | 0.70 | 2,444,070 | 0.007 |
|  | 0.05-0.1 | 0.31 | 0.69 | 1,180,341 | 0.016 | 0.22 | 0.78 | 1,097,724 | 0.011 |
|  | 0.1-0.2 | 0.31 | 0.69 | 1,379,163 | 0.023 | 0.21 | 0.79 | 1,287,160 | 0.015 |
|  | 0.2-0.3 | 0.31 | 0.69 | 913,809 | 0.028 | 0.21 | 0.79 | 853,249 | 0.018 |
|  | 0.3-0.4 | 0.30 | 0.70 | 698,053 | 0.031 | 0.20 | 0.80 | 652,017 | 0.020 |
|  | 0.4-0.5 | 0.31 | 0.69 | 555,041 | 0.033 | 0.21 | 0.79 | 518,228 | 0.021 |
|  | 0.5-0.6 | 0.31 | 0.69 | 460,852 | 0.033 | 0.21 | 0.79 | 430,399 | 0.021 |
|  | 0.6-0.7 | 0.31 | 0.69 | 396,445 | 0.032 | 0.21 | 0.79 | 369,703 | 0.021 |
|  | 0.7-0.8 | 0.32 | 0.68 | 329,217 | 0.029 | 0.21 | 0.79 | 306,753 | 0.019 |
|  | 0.8-0.9 | 0.32 | 0.68 | 272,693 | 0.024 | 0.21 | 0.79 | 254,014 | 0.016 |
|  | 0.9-1 | 0.32 | 0.68 | 391,595 | 0.013 | 0.14 | 0.86 | 366,541 | 0.008 |

\*Number of variants
\*\*Mean confidence interval lengths
Abbreviations: AF Allele frequency; GAFA Global -Ancestral Frequency estimation in Admixed populations;
LAFA Local -Ancestral Frequency estimation in Admixed populations.

**Figure 2:**
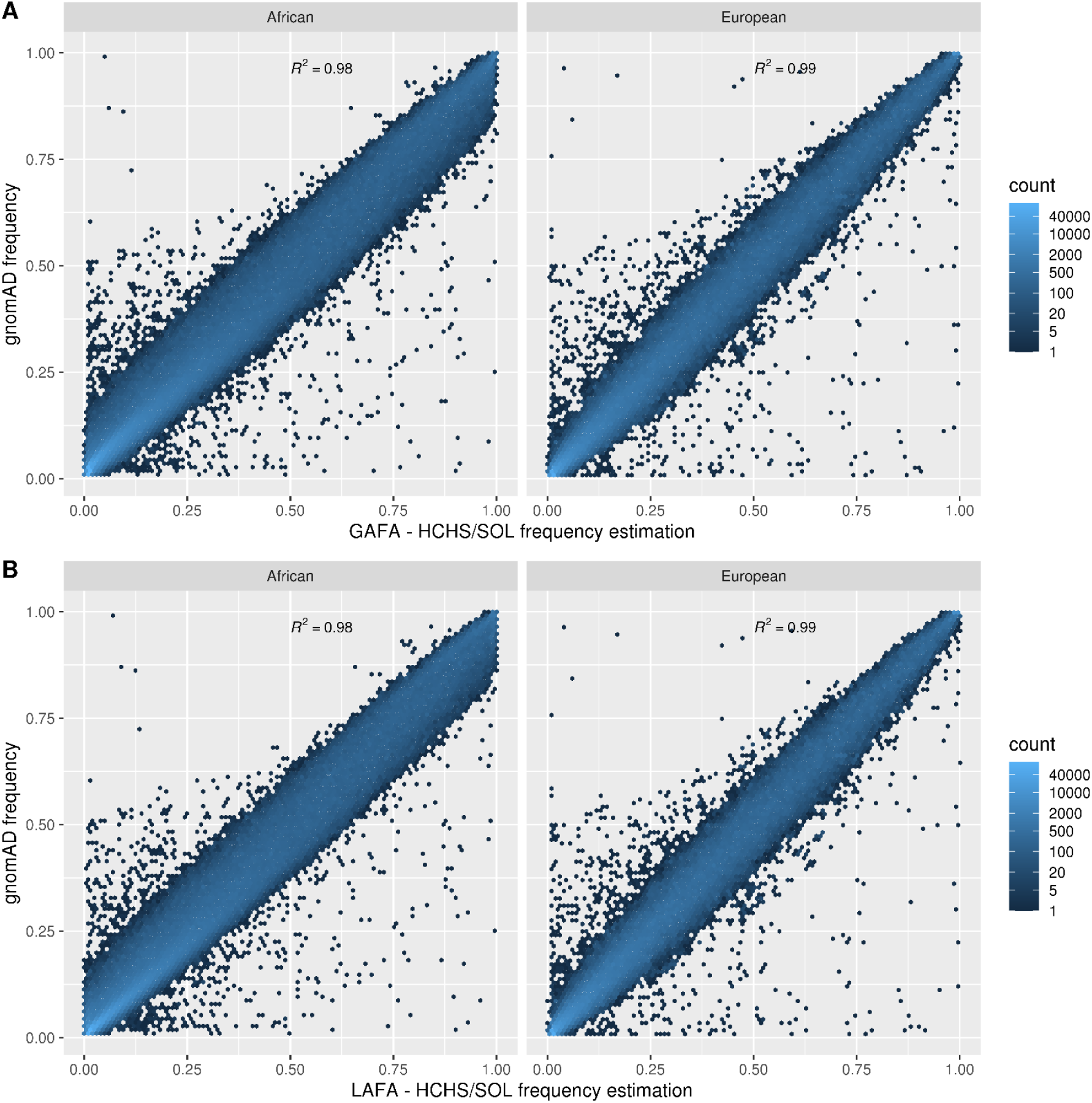
Scatter plots of estimated ancestry-specific allele frequencies in HCHS/SOL chromosome 2 to corresponding gnomAD non-Finnish European and African frequencies respectively (A) using GAFA (no. variants: African=1,239,958 European=819,710) (B) using LAFA (no. variants: African=1,168,271 European= 775,749).

### Correlation of estimated ancestry-specific allele frequencies between the GAFA and LAFA for each of the 3 ancestries

Figure 3 presents strong correlations of the chromosome 2 estimated ancestry-specific allele frequencies in the HCHS/SOL population between GAFA and LAFA for each of the three ancestral populations. The European’s correlation is stronger than the Africans and Amerindians. This is probably due to their larger effective sample size in the HCHS/SOL, enabling a more precise estimation of the alleles’ frequencies (effn based on global proportion ancestries: African=1,296 European=4,912 Amerindian=2,725). All other chromosomes’ correlations are presented in Supplementary Figure 4.

**Figure 3:**
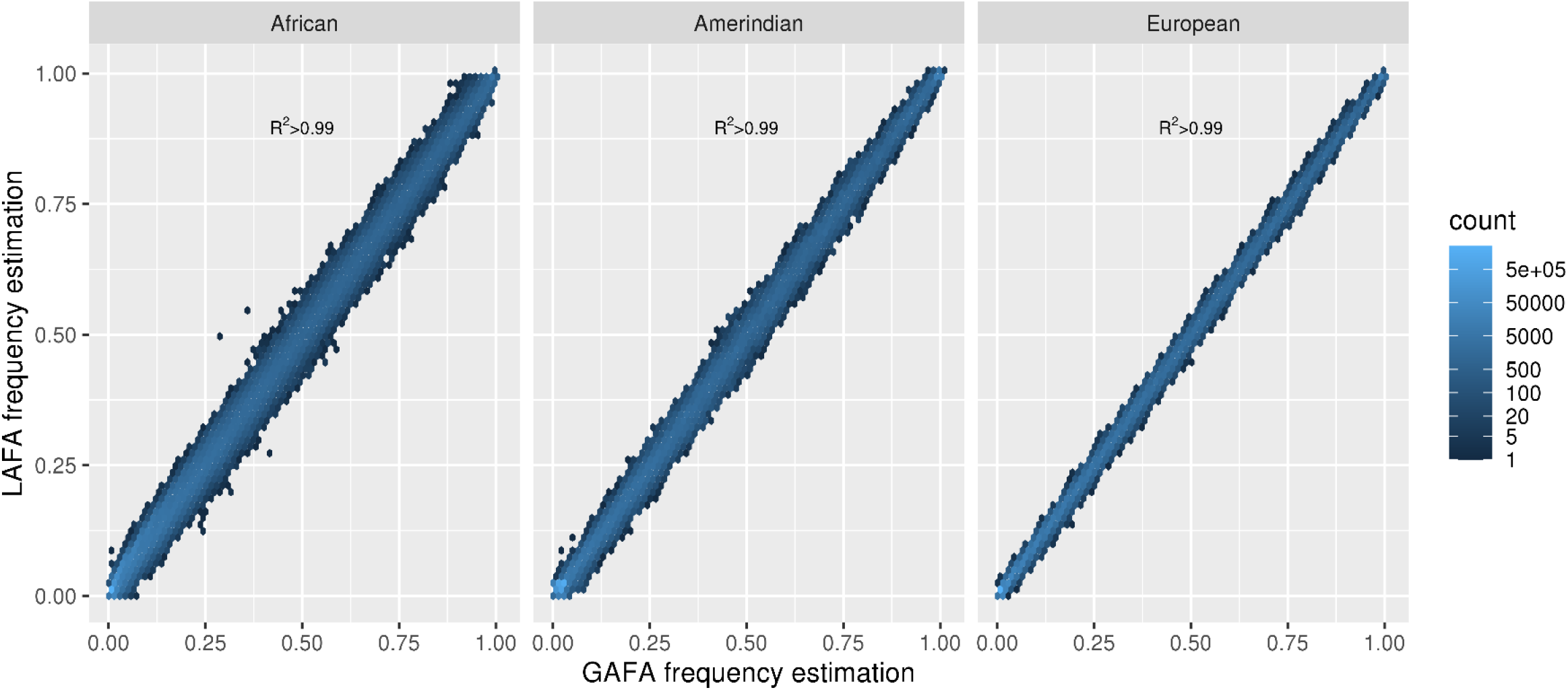
Scatter plots of the estimated ancestry-specific allele frequencies in chromosome 2 in the HCHS/SOL population between GAFA and LAFA for each of the three ancestral populations (no. variants=9,308,589).

**Figure 4:**
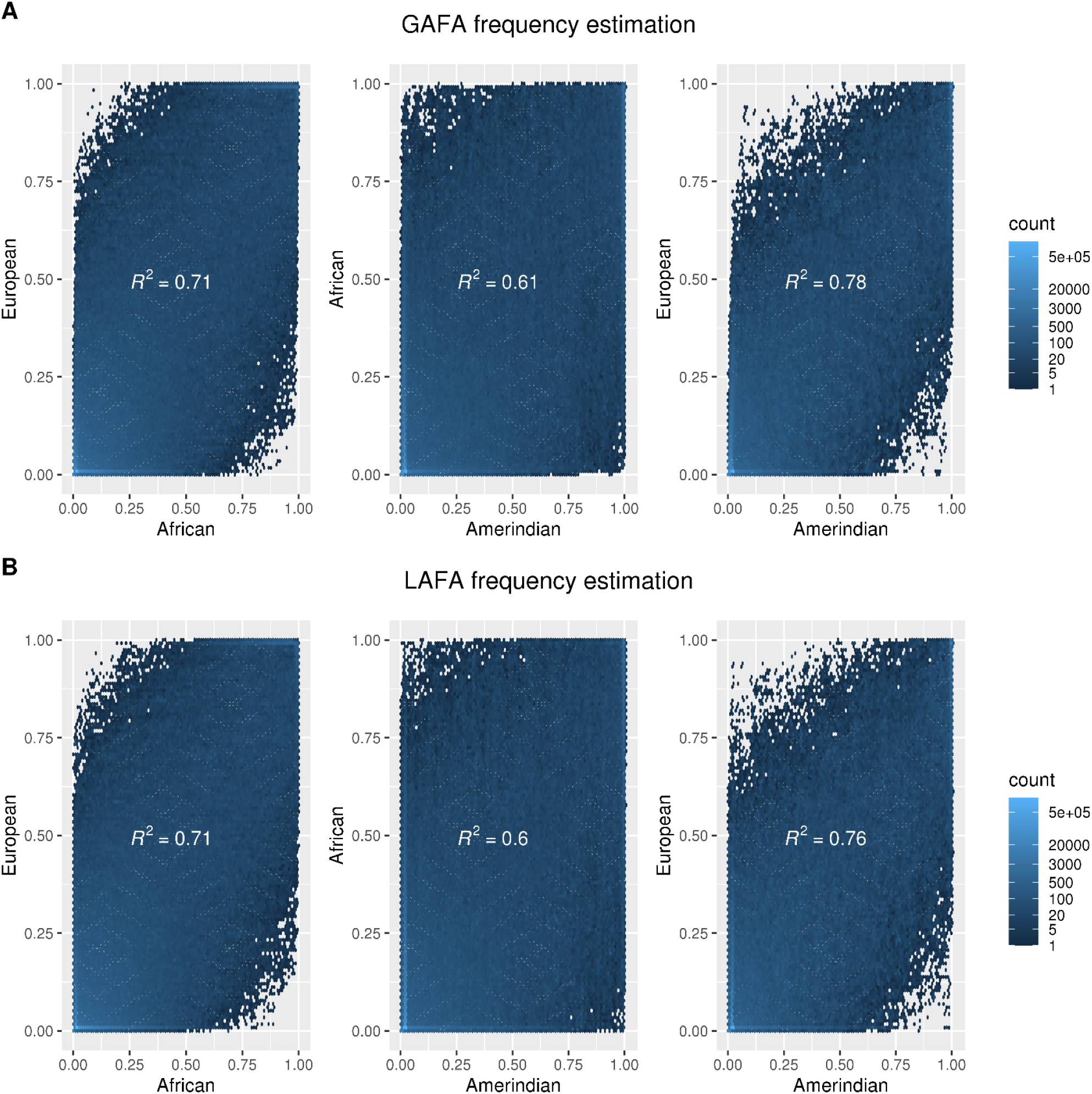
Scatter plots of the estimated ancestry-specific allele frequencies in chromosome 2 in the HCHS/SOL population between the three ancestral populations, for (A) GAFA (B) LAFA.

### Correlation of the estimated ancestry-specific allele frequencies between different ancestries

Figure 4 presents weak correlations of the estimated ancestry-specific allele frequencies for chromosome 2 variants in the HCHS/SOL population between the three ancestral populations, for both GAFA and LAFA. The squared Pearson correlation coefficient is strongest when comparing Amerindian to European ancestral frequencies (GAFA: R^2^=0.78, LAFA: R^2^=0.76), followed by the comparison of African to European (GAFA: R^2^=0.71, LAFA: R^2^=0.71), and weakest in the comparison of African to Amerindian (GAFA: R^2^=0.61, LAFA: R^2^=0.6). Similar correlations of all other chromosomes are presented in Supplementary Figures 5 (GAFA) and 6 (LAFA).

### Evaluating the algorithm convergence rate of GAFA and LAFA by frequency boundary conditions

Summary statistics of HCHS/SOL alleles calculated using AFA vs. alleles that failed calculation are presented in Supplementary Table 3. For variants on chromosome 2, 92.3% were calculated using GAFA (n=3,299310,366) and 88.8% were calculated using LAFA (n=3,175,914). Low MAF is likely the main reason for failed ancestral MAF calculation in admixed populations using our method. LAFA’s successful calculation percentage is lower compared to GAFA since the LAIs do not encompass the whole genome, and the liftover from GRCh38 to GRCh37 (in order to match each variant to its LAI) also failed for some variants. The number of overlapped calculated variants in both methods on chromosome 2 is n=3,102,863, while n=192,993 variants were successfully calculated only in GAFA and n=70,641 variants were successfully calculated only in LAFA. This emphasizes the importance of developing both methods and their potential to complement each other.

## Discussion

We developed a computationally efficient method for estimating ancestry-specific variant frequencies in admixed populations (AFA) based on either the rather widely available global proportion ancestry (GAFA) or LAIs (LAFA). Simulations have shown high accuracy of the estimated frequencies for both options, with increasing accuracy dependent on ancestral effective population and MAF, and with a slight advantage for LAFA over GAFA. We applied our method to the admixed Hispanic/Latinos population from HCHS/SOL with three predominant continental ancestries: European, African, and Amerindian, and demonstrated speed, simplicity of calculation, and a highly successful frequency estimation rate.

Comparison of the European and African estimated ancestral specific frequencies to the respective gnomAD frequencies demonstrated strong positive correlations. We did not expect perfect correlation with the respective gnomAD frequencies, since evolutionary forces such as genetic drift, mutagenesis, and natural selection are expected to accumulate and result in frequency differences. The correlation found in Europeans is somewhat stronger compared to the Africans. This is likely due to two reasons: first, individuals of African ancestries are characterized by a greater level of genetic diversity compared to Europeans^25^, so allele frequency comparisons between two populations of African ancestral origin will demonstrate a larger difference compared to frequency comparisons between two populations of European ancestral origin. Second, the effective sample size of European ancestry in the HCHS/SOL was substantially larger than the African effective sample size, enabling a more precise estimation of allele frequencies.

We provide a genome-wide dataset of U.S. Hispanic/Latino ancestry-specific allele frequencies estimated based on the HCHS/SOL for all variants with a frequency between 5%-95% in at least one of the three ancestral populations, using GAFA (n= 9,808,089) and LAFA (n= 9,844,093). To our knowledge, this is the first published genome-wide dataset of ancestral frequencies in an admixed population. Specifically, the Amerindian allele frequency estimation is otherwise unavailable. Inter-HCHS/SOL ancestral frequencies present the strongest correlations between Amerindians and Europeans, followed by the Africans and Europeans followed by Africans and Amerindians. These findings agree with the dominant paleoanthropology hypothesis of the African origin of modern humans, followed by migration to Europe, followed by other migrations to Asia and America^26^. Stronger bottlenecks (founder effect) in Amerindians led to more drifts and hence more differences in Amerindian compared to African frequencies. Thus, our dataset can serve as a unique resource for genetic epidemiology studies supporting research of personalized health in admixed populations.

The advantages of our method are the ability to estimate ancestral frequencies and CIs of genotyped or imputed variants in admixed populations with an unlimited number of ancestries, with no need for phased data, on a genome-wide scale. The algorithm is applicable for phased data as well. Thus, our method is simple, effective, and enables a wider usage. It can be applied using either global proportions of genetic ancestries (GAFA) or LAI proportions encompassing the variant (LAFA). GAFA is a computationally simpler process compared to LAFA, and it encompasses all regions of the genome. However, it assumes a uniform distribution of ancestries throughout the genome; which is slightly less precise. Comparison of both GAFA and LAFA shows strong correlations for variants calculated by both methods and shows some variants could be calculated by using only one of the methods, complementing each other and emphasizing the advantage of using both options. Specifically, LAFA is more precise; but the algorithm may not converge when using LAFA so that frequency estimates were not obtained, while GAFA may converge for these variants. We think that this is likely due to local ancestry inference errors: when using LAFA, the ancestral probabilities assigned by the algorithm at the segment take values *p*_*i*1_, … , *p*_*iK*_ ∈ {0,0.5,1}. Thus, if in all LAIs from a specific ancestry the observed MAC is 0, it may lead to non-convergence. Non-convergence may also arise from a lack of HWE in LAIs from a certain ancestry. Depending on effective population sample sizes, our method may perform less well for low MAFs variants. First, estimation depends on the effective sample sizes of the ancestral origins and the ancestry-specific frequencies (e.g. having enough counts). Second, AFA methods apply maximum likelihood estimation of Binomial likelihoods, which cannot be evaluated by the optimization algorithm at the boundaries of the parameter space (frequencies of 0 or 1; though the likelihood is computed at the boundary). Therefore, very few minor allele counts in one of the genetic ancestries may lead to non-convergence of the algorithm, unless box constraints are placed (e.g., limiting the frequencies to be estimated within the interval [0.01, 0.99]), so that frequencies outside the interval cannot be estimated.

## Acknowledgments

The authors thank the staff and participants of HCHS/SOL for their important contributions. Investigator’s website - http://www.cscc.unc.edu/hchs/. The Hispanic Community Health Study/Study of Latinos is a collaborative study supported by contracts from the National Heart, Lung, and Blood Institute (NHLBI) to the University of North Carolina (HHSN268201300001I / N01-HC-65233), University of Miami (HHSN268201300004I / N01-HC-65234), Albert Einstein College of Medicine (HHSN268201300002I / N01-HC-65235), University of Illinois at Chicago (HHSN268201300003I / N01-HC-65236 Northwestern Univ), and San Diego State University (HHSN268201300005I / N01-HC-65237). The following Institutes/Centers/Offices have contributed to the HCHS/SOL through a transfer of funds to the NHLBI: National Institute on Minority Health and Health Disparities, National Institute on Deafness and Other Communication Disorders, National Institute of Dental and Craniofacial Research, National Institute of Diabetes and Digestive and Kidney Diseases, National Institute of Neurological Disorders and Stroke, NIH Institution-Office of Dietary Supplements. The Genetic Analysis Center at the University of Washington was supported by NHLBI and NIDCR contracts (HHSN268201300005C AM03 and MOD03).

The authors would also like to acknowledge the use of the Trans-Omics in Precision Medicine (TOPMed) program imputation panel (freeze-8 version) supported by the National Heart, Lung and Blood Institute (NHLBI); see www.nhlbiwgs.org. TOPMed study investigators contributed data to the reference panel, which was accessed through https://imputation.biodatacatalyst.nhlbi.nih.gov. The panel was constructed and implemented by the TOPMed Informatics Research Center at the University of Michigan (3R01HL-117626-02S1; contract HHSN268201800002I). The TOPMed Data Coordinating Center (R01HL-120393; U01HL-120393; contract HHSN268201800001I) provided additional data management, sample identity checks, and overall program coordination and support. We gratefully acknowledge the studies and participants who provided biological samples and data for TOPMed.

## Author contributions

E.G.H. and T.S. conceived the presented method and drafted the manuscript. T.S. developed the statistical method and E.G.H. performed the computations. Q.S. performed the imputation analysis. All authors critically reviewed the manuscript.

