## Supplementary material for "AFA: Computationally efficient Ancestral Frequency estimation in Admixed populations: the Hispanic Community Health Study/Study of Latinos": 20210806_Supplementary.docx

**Supplementary Figure 1:** Results from simulations estimating ancestry-specific frequencies of a bi-allelic variant in a homogenous population (single ancestry), based on AFA (Global and Local -Ancestral Frequency estimation in Admixed populations are identical for non-admixed populations). Various settings include a different effective sample size of effn (x-axis) and different expected minor allele frequencies (indicated in the upper title of each graph). We performed 1,000 simulation replicates of each scenario. Each dot represents the mean frequency of 1,000 simulation replicates each line represents the 95% interval of the simulation replicates.

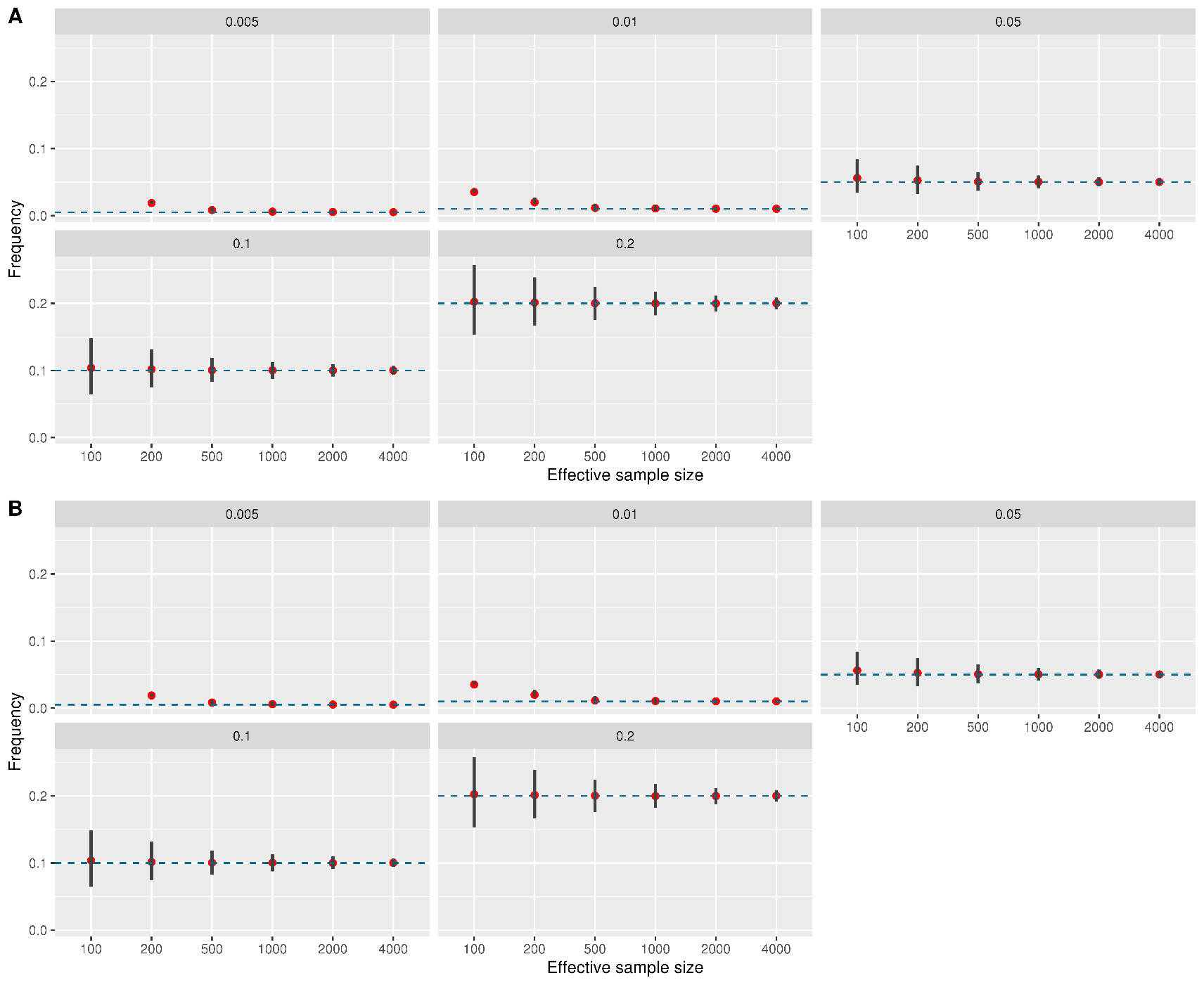
**Supplementary Figure 2:** Scatter plots of GAFA estimated ancestry-specific allele frequencies in the HCHS/SOL population to corresponding gnomAD non-Finnish European and African frequencies
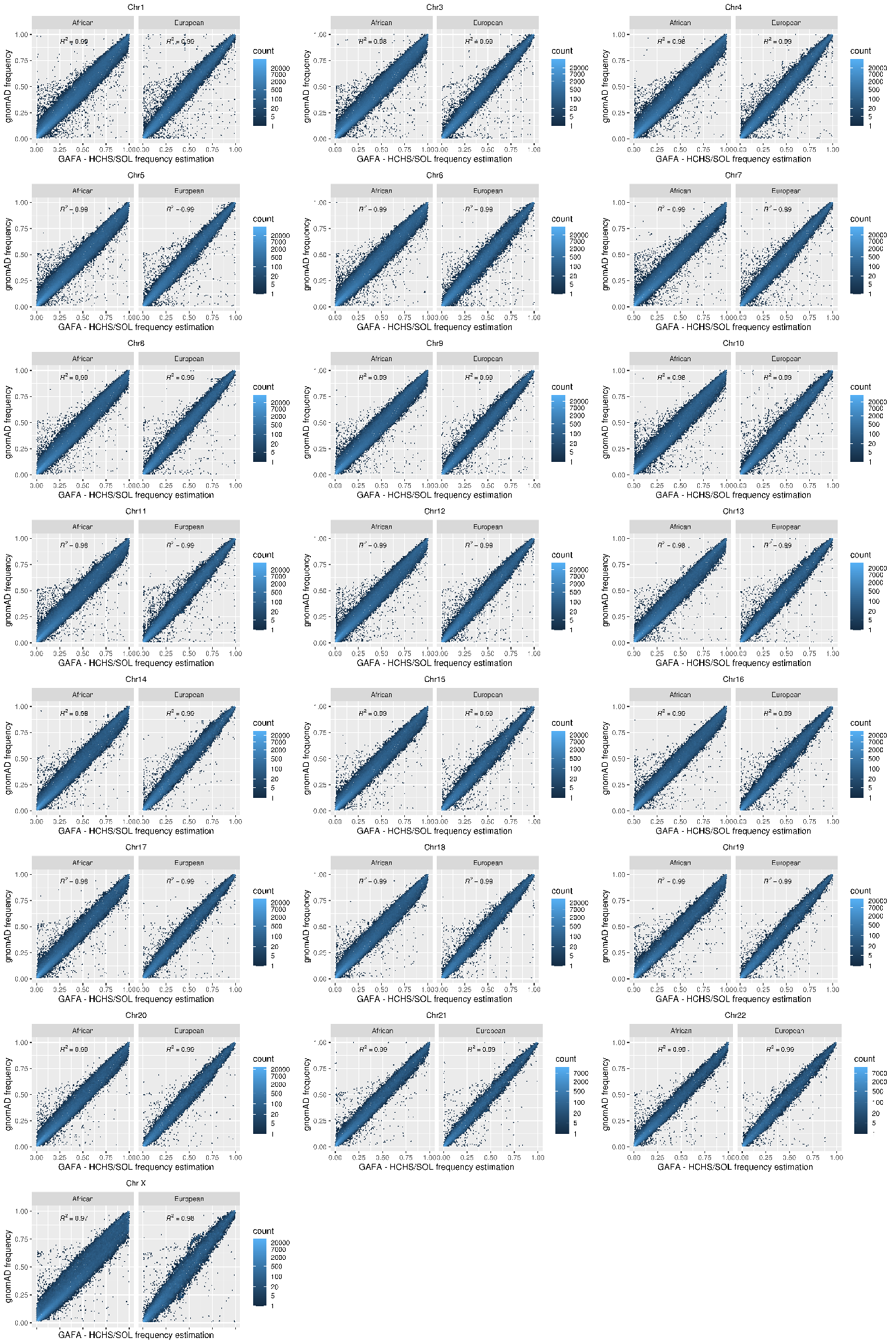
respectively for each chromosome.

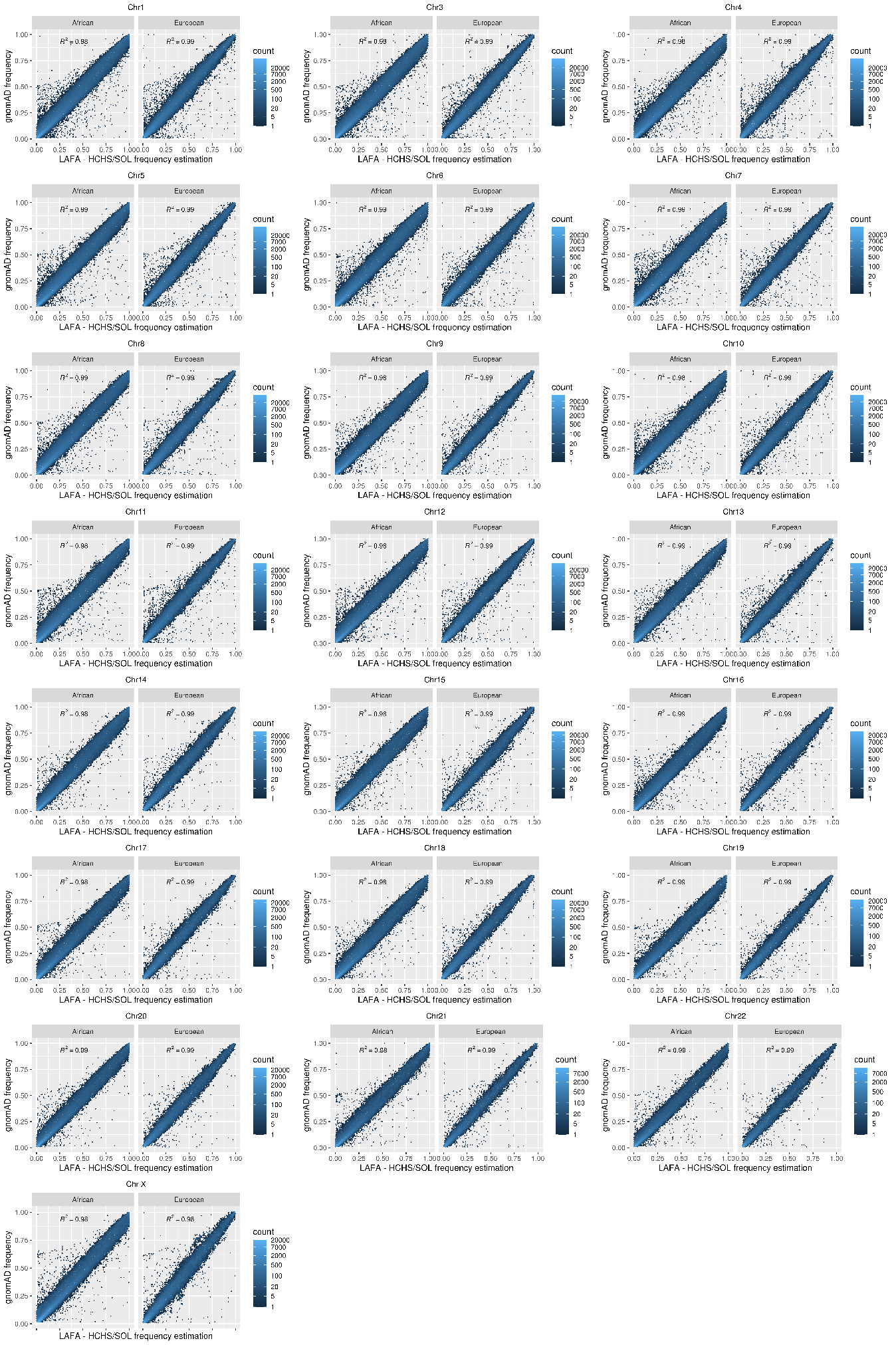
**Supplementary Figure 3:** Scatter plots of LAFA estimated ancestry-specific allele frequencies in the HCHS/SOL population to corresponding gnomAD non-Finnish European and African frequencies respectively for each chromosome.

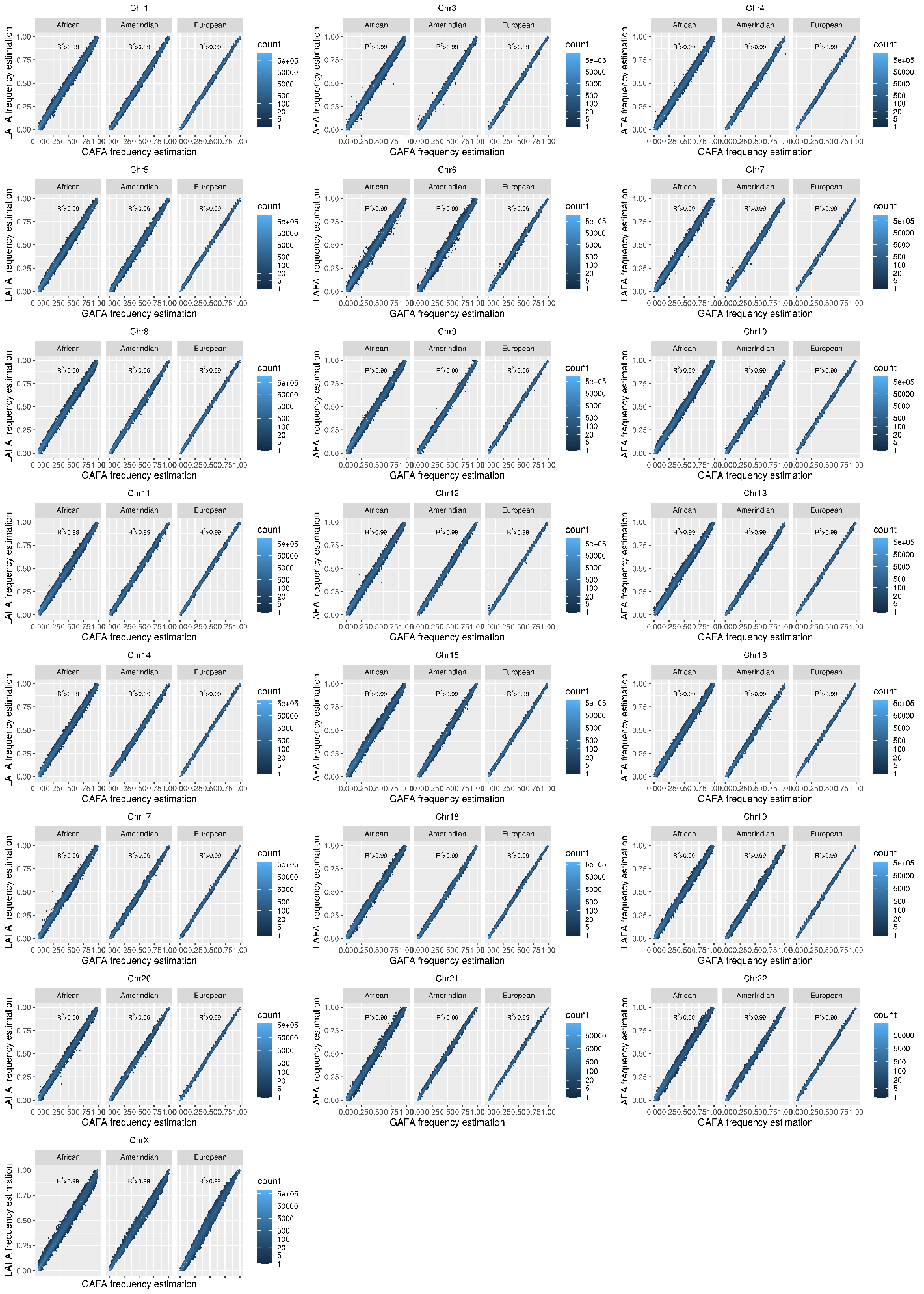
**Supplementary Figure 4:** Scatter plots of the estimated ancestry-specific allele frequencies in the HCHS/SOL population between GAFA and LAFA for each of the three ancestral populations for each chromosome.

**Supplementary Figure 5:** Scatter plots of GAFA estimated ancestry-specific allele frequencies in the HCHS/SOL population for all chromosomes, between the three ancestral populations.

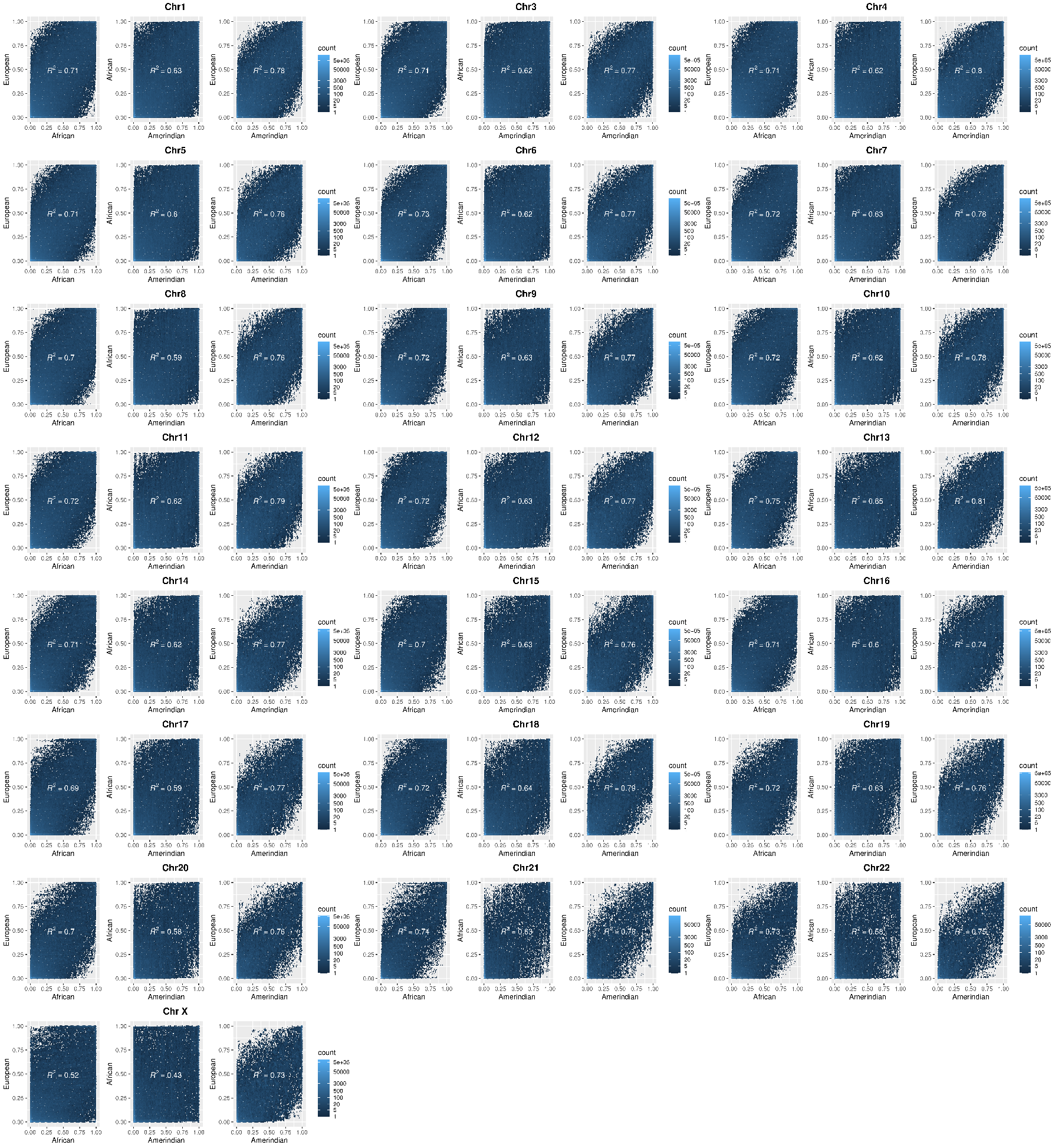

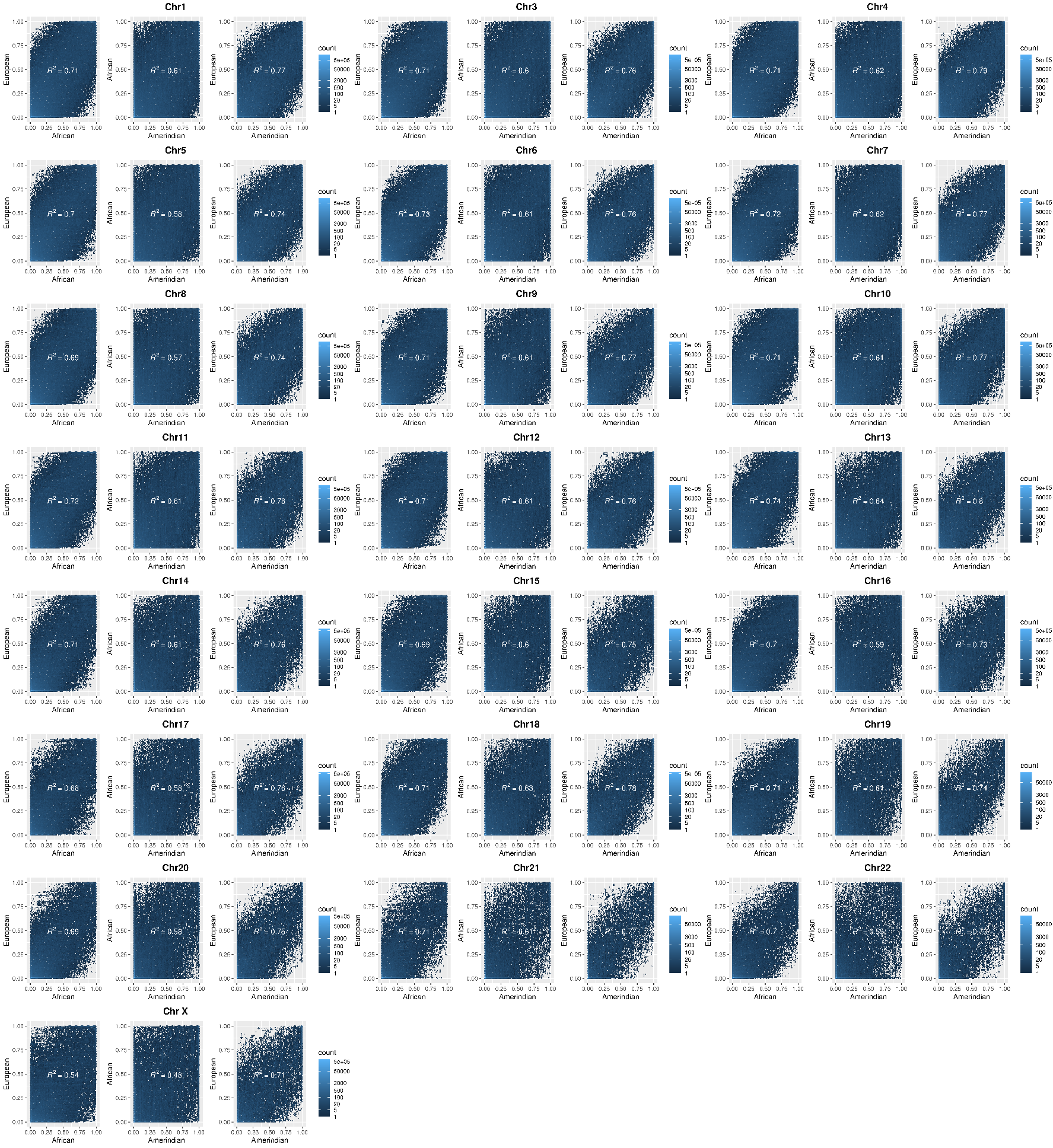
**Supplementary Figure 6:** Scatter plots of LAFA estimated ancestry-specific allele frequencies in the HCHS/SOL population for all chromosomes, between the three ancestral populations.

**Supplementary Table 1**: Results from simulation studies of frequency estimation of a bi-allelic variant in a homogenous population (single ancestry), using the AFA (Ancestral Frequency estimation in Admixed populations) algorithm, by different effective sample sizes and different expected minor allele frequencies. For each of the settings, we tested 1,000 simulation replicates and calculated the mean frequency estimate, the difference, and ratio of the mean observed frequency and the expected frequency, the RMSE, the percentage of CI including the expected frequency (coverage), and the 95% interval of the observed frequency.

|  |  | Single ancestry population | | | | | | |
| --- | --- | --- | --- | --- | --- | --- | --- | --- |
| Expected MAF | Effective N | MAF Mean | MAF difference | MAF ratio | RMSE | CI % coverage | interval 2.5% | interval 97.5% |
| 0.005 | 100 | - | - | - | - | - | - | - |
|  | 200 | 0.019 | 0.014 | 3.770 | 0.014 | 0.5 | 0.017 | 0.022 |
|  | 500 | 0.008 | 0.003 | 1.664 | 0.004 | 0.97 | 0.007 | 0.012 |
|  | 1000 | 0.006 | 0.001 | 1.173 | 0.002 | 0.97 | 0.004 | 0.010 |
|  | 2000 | 0.005 | 0.000 | 1.076 | 0.001 | 0.95 | 0.003 | 0.008 |
|  | 4000 | 0.005 | 0.000 | 1.034 | 0.001 | 0.94 | 0.004 | 0.007 |
| 0.01 | 100 | 0.035 | 0.025 | 3.521 | 0.025 | 0.89 | 0.035 | 0.040 |
|  | 200 | 0.020 | 0.010 | 1.988 | 0.010 | 0.97 | 0.017 | 0.027 |
|  | 500 | 0.011 | 0.001 | 1.142 | 0.003 | 0.98 | 0.007 | 0.018 |
|  | 1000 | 0.011 | 0.001 | 1.057 | 0.002 | 0.94 | 0.007 | 0.015 |
|  | 2000 | 0.010 | 0.000 | 1.025 | 0.002 | 0.96 | 0.008 | 0.013 |
|  | 4000 | 0.010 | 0.000 | 1.015 | 0.001 | 0.95 | 0.008 | 0.012 |
| 0.05 | 100 | 0.056 | 0.006 | 1.121 | 0.015 | 0.99 | 0.035 | 0.084 |
|  | 200 | 0.052 | 0.002 | 1.049 | 0.011 | 0.95 | 0.032 | 0.075 |
|  | 500 | 0.051 | 0.001 | 1.014 | 0.007 | 0.94 | 0.037 | 0.065 |
|  | 1000 | 0.050 | 0.000 | 1.008 | 0.005 | 0.95 | 0.041 | 0.060 |
|  | 2000 | 0.050 | 0.000 | 1.003 | 0.003 | 0.95 | 0.044 | 0.057 |
|  | 4000 | 0.050 | 0.000 | 1.005 | 0.002 | 0.95 | 0.045 | 0.055 |
| 0.1 | 100 | 0.104 | 0.004 | 1.037 | 0.021 | 0.95 | 0.064 | 0.149 |
|  | 200 | 0.102 | 0.002 | 1.016 | 0.015 | 0.96 | 0.075 | 0.132 |
|  | 500 | 0.100 | 0.000 | 1.004 | 0.009 | 0.95 | 0.083 | 0.119 |
|  | 1000 | 0.100 | 0.000 | 1.004 | 0.007 | 0.95 | 0.087 | 0.113 |
|  | 2000 | 0.100 | 0.000 | 0.999 | 0.005 | 0.95 | 0.091 | 0.110 |
|  | 4000 | 0.100 | 0.000 | 1.002 | 0.003 | 0.95 | 0.094 | 0.107 |
| 0.2 | 100 | 0.202 | 0.002 | 1.012 | 0.028 | 0.95 | 0.153 | 0.257 |
|  | 200 | 0.201 | 0.001 | 1.006 | 0.019 | 0.97 | 0.167 | 0.239 |
|  | 500 | 0.200 | 0.000 | 1.001 | 0.013 | 0.95 | 0.176 | 0.225 |
|  | 1000 | 0.200 | 0.000 | 0.999 | 0.009 | 0.94 | 0.182 | 0.218 |
|  | 2000 | 0.200 | 0.000 | 0.999 | 0.006 | 0.96 | 0.188 | 0.212 |
|  | 4000 | 0.200 | 0.000 | 1.001 | 0.004 | 0.95 | 0.191 | 0.209 |

Abbreviations: *MAF* minor allele frequency; *CI* confidence interval; *RMSE* root mean squared error.

| **Supplementary Table 2:** Number of the total reported estimated variant frequencies per chromosome in HCHS/SOL, stratified by boundary condition, calculated via GAFA or LAFA. | | | | | | |
| --- | --- | --- | --- | --- | --- | --- |
|  | GAFA | | | LAFA | | |
| Chr. | Total | Boundary 1E-05 (%) | Boundary 1E-02 | Total | Boundary 1E-05 (%) | Boundary 1E-02 |
| 1 | 3,015,926 | 693,503 (22.99) | 2,322,423 | 2,860,144 | 445,290 (15.57) | 2,414,854 |
| 2 | 3,295,856 | 751,537 (22.8) | 2,544,319 | 3,173,504 | 481,129 (15.16) | 2,692,375 |
| 3 | 2,743,144 | 643,966 (23.48) | 2,099,178 | 2,634,244 | 417,617 (15.85) | 2,216,627 |
| 4 | 2,752,093 | 661,779 (24.05) | 2,090,314 | 2,632,654 | 438,011 (16.64) | 2,194,643 |
| 5 | 2,519,640 | 585,336 (23.23) | 1,934,304 | 2,420,555 | 368,344 (15.22) | 2,052,211 |
| 6 | 2,411,777 | 612,582 (25.4) | 1,799,195 | 2,313,271 | 379,800 (16.42) | 1,933,471 |
| 7 | 2,242,807 | 540,367 (24.09) | 1,702,440 | 2,114,380 | 344,519 (16.29) | 1,769,861 |
| 8 | 2,165,759 | 496,885 (22.94) | 1,668,874 | 2,091,712 | 315,476 (15.08) | 1,776,236 |
| 9 | 1,681,237 | 396,484 (23.58) | 1,284,753 | 1,608,965 | 250,008 (15.54) | 1,358,957 |
| 10 | 1,907,281 | 465,530 (24.41) | 1,441,751 | 1,808,938 | 296,368 (16.38) | 1,512,570 |
| 11 | 1,886,493 | 446,785 (23.68) | 1,439,708 | 1,817,103 | 289,395 (15.93) | 1,527,708 |
| 12 | 1,840,735 | 437,086 (23.75) | 1,403,649 | 1,737,222 | 275,193 (15.84) | 1,462,029 |
| 13 | 1,376,312 | 339,010 (24.63) | 1,037,302 | 1,326,019 | 223,561 (16.86) | 1,102,458 |
| 14 | 1,238,478 | 298,009 (24.06) | 940,469 | 1,175,568 | 184,692 (15.71) | 990,876 |
| 15 | 1,115,118 | 261,894 (23.49) | 853,224 | 1,060,977 | 167,129 (15.75) | 893,848 |
| 16 | 1,191,194 | 268,591 (22.55) | 922,603 | 1,098,816 | 169,806 (15.45) | 929,010 |
| 17 | 1,087,574 | 247,678 (22.77) | 839,896 | 999,265 | 151,927 (15.2) | 847,338 |
| 18 | 1,082,746 | 261,325 (24.14) | 821,421 | 1,033,775 | 170,768 (16.52) | 863,007 |
| 19 | 864,582 | 211,835 (24.5) | 652,747 | 768,048 | 124,533 (16.21) | 643,515 |
| 20 | 863,716 | 203,289 (23.54) | 660,427 | 814,127 | 127,206 (15.62) | 686,921 |
| 21 | 509,043 | 123,941 (24.35) | 385,102 | 480,467 | 80,656 (16.79) | 399,811 |
| 22 | 521,404 | 126,553 (24.27) | 394,851 | 474,338 | 74,521 (15.71) | 399,817 |
| X | 1,435,812 | 278,315 (19.38) | 1,157,497 | 1,247,194 | 157,811 (12.65) | 1,089,383 |
| Total | 39,748,727 | 9,352,280 (23.53) | 30,396,447 | 37,691,286 | 5,933,760 (15.74) | 31,757,526 |
| Abbreviations: GAFA Global -Ancestral Frequency estimation in Admixed populations; LAFA Local -Ancestral Frequency estimation in Admixed populations. | | | | | | |

| **Supplementary Table 3**: Summary statistics of chromosome 2 ancestry-specific calculated alleles in HCHS/SOL using AFA vs. non-calculated alleles. | | | | | | | | |
| --- | --- | --- | --- | --- | --- | --- | --- | --- |
|  | | | Genotype | | | Allele Frequency | HWE  P-val | R2 (imputation quality) |
|  |  |  | nAA * | nAB | nBB |  |  |  |
| GAFA | Calculated n=3,299,101 (92.7%) | Mean | 10818.21 | 702.98 | 406.81 | 0.0636 | 0.678 | 0.96 |
|  |  | SD | 2646.993 | 1485.97 | 1560.10 | 0.1712 | 0.429 | 0.06 |
|  |  | Median | 11884 | 44 | 0 | 0.0018 | 1 | 0.99 |
|  |  | Min | 0 | 2 | 0 | 0.0003 | 0 | 0.60 |
|  |  | Max | 11924 | 6181 | 11921 | 0.9997 | 1 | 1 |
|  | Non-Calculated n=259,943 | Mean | 11921.15 | 6.83 | 0.02 | 0.0003 | 0.980 | 0.94 |
|  |  | SD | 1.220091 | 1.24 | 0.15 | 0.0001 | 0.139 | 0.07 |
|  |  | Median | 11922 | 6 | 0 | 0.0003 | 1 | 0.97 |
|  |  | Min | 11910 | 2 | 0 | 0.0003 | 1.23E-16 | 0.60 |
|  |  | Max | 11924 | 18 | 5 | 0.0008 | 1 | 1 |
| LAFA | Calculated n=3,175,727 (89.2%) | Mean | 10840.08 | 688.23 | 399.69 | 0.0624 | 0.685 | 0.96 |
|  |  | SD | 2627.187 | 1473.81 | 1549.16 | 0.1699 | 0.427 | 0.06 |
|  |  | Median | 11887 | 41 | 0 | 0.0017 | 1 | 0.99 |
|  |  | Min | 0 | 2 | 0 | 0.0003 | 0 | 0.60 |
|  |  | Max | 11924 | 6181 | 11921 | 0.9997 | 1 | 1 |
|  | Non-Calculated n=383,317 | Mean | 11385.01 | 353.10 | 189.89 | 0.0307 | 0.822 | 0.95 |
|  |  | SD | 1902.704 | 1101.52 | 1063.08 | 0.1207 | 0.360 | 0.07 |
|  |  | Median | 11920 | 8 | 0 | 0.0003 | 1 | 0.98 |
|  |  | Min | 0 | 2 | 0 | 0.0003 | 1.44E-238 | 0.60 |
|  |  | Max | 11924 | 6088 | 11892 | 0.9985 | 1 | 1 |
| **Abbreviations:** *AFA* Ancestral Frequency estimation in Admixed populations, *GAFA* Global -Ancestral Frequency estimation in Admixed populations, *LAFA* Local -Ancestral Frequency estimation in Admixed populations, *HWE* Hardy Weinberg Equilibrium, *SD* standard deviation.  *A is the common allele. | | | | | | | | |
